# Should deep-sequenced amplicons become the new gold-standard for analysing malaria drug clinical trials?

**DOI:** 10.1101/2021.03.23.436602

**Authors:** Sam Jones, Katherine Kay, Eva Maria Hodel, Maria Gruenberg, Anita Lerch, Ingrid Felger, Ian Hastings

## Abstract

**Background:** Regulatory clinical trials are required to ensure the continued supply and deployment of effective antimalarial drugs. Patient follow-up in such trials typically lasts several weeks as the drugs have long half-lives and new infections often occur during this period. “Molecular correction” is therefore used to distinguish drug failures from new infections. The current WHO-recommend method for molecular correction uses length-polymorphic alleles at highly diverse loci but is inherently poor at detecting low density clones in polyclonal infections. This likely leads to substantial underestimates of failure rates, delaying the replacement of failing drugs with potentially lethal consequences. Deep sequenced amplicons (AmpSeq) substantially increase the detectability of low-density clones and may offer a new “gold standard” for molecular correction.

**Methods:** Pharmacological simulation of clinical trials was used to evaluate the suitability of AmpSeq for molecular correction. We investigated the impact of factors such as the number of amplicon loci analysed, the informatics criteria used to distinguish genotyping ‘noise’ from real low density signals, the local epidemiology of malaria transmission, and the potential impact of genetic signals from gametocytes.

**Results:** AmpSeq greatly improved molecular correction and provided accurate drug failure rate estimates. The use of 3 to 5 amplicons was sufficient, and simple, non-statistical, criteria could be used to classify recurrent infections as drug failures or new infections.

**Conclusions:** These results strongly endorse the deployment of AmpSeq as the standard for molecular correction in regulatory trials, with its potential extension into routine surveillance once the requisite technical support becomes established.

## 1. Introduction

Clinical trials and Therapeutic efficacy studies (TES) of anti-malarial drugs are key components of public health provision in malaria endemic countries. The role of clinical trials is to ensure a steady supply of effective drugs, while the role of TES is to provide ongoing surveillance on the efficacy of local front-line drugs, and enable rapid replacement of failing drugs to avoid the increased morbidity and mortality associated with drug resistance (1).

In principle clinical trials and TES are simple: patients are treated with the drug being evaluated, and the number of patients with drug failures are counted to provide estimates of drug effectiveness. In practice, this is challenging because most current malaria drugs have long half-lives such that drug failures are suppressed and may only become patent several weeks after treatment. Ongoing malaria transmission means that patients often acquire new infections during the long follow-up period so a key methodological requirement of clinical trials and TES is to correctly classify patients returning with detectable malaria parasites during follow-up (termed “recurrences”) as either drug failures (termed “recrudescences”) or new infections (sometimes termed “reinfection”). This classification is achieved using molecular correction protocols which rely on genotyping malaria parasites in the blood of the infected patient at the time of rug treatment and if that patient returns with a recurrence at any time during the 4-6 week follow-up period (2). ‘Matching’ alleles between treatment and follow-up samples indicate a drug failure, while ‘mis-matched’ allele(s) indicate a new infection. Deciding whether or not the samples match is highly problematic, and the number of shared alleles required to define a match (and hence a drug failure) depends on the methodology used (3–5).

The World Health Organization (WHO) recommends three markers for molecular correction (2) i.e. merozoite surface protein-1 (*msp-1*), merozoite surface protein-2 (*msp-2*) and glutamate rich protein (*glurp*) though alternative markers are available such as the microsatellite markers used by the Centers for Disease Control and Prevention (6). The three WHO markers and the CDC microsatellite markers all rely on identifying alleles by their lengths (in contrast to amplicons, discussed later, where allele differ in their sequence). Length-polymorphic genotyping uses PCR amplification of molecular markers in patient blood samples, followed by fragment sizing of the PCR products using agarose gels or capillary electrophoresis. Malaria infections often contain multiple parasite clones both at treatment (when the multiplicity of infection, MOI, is typically 3 to 8 malaria clones per person in high transmission areas) and at recurrence (where patients may present with a mixture of recrudescence clones and new infections). There is substantial variation is the density of individual clones in these multi-clonal infections and existing methods based on length polymorphism are notoriously poor at detecting low-density clones. These methods typically regard genetic signals less than round 20% to 30% of the major signal as ‘noise’, meaning any clones whose density is less than 20% to 30% that of the dominant clone are not identified. There have been recent calls for a review of the WHO-recommended methodology, (5, 7) following scientific discussion of misclassification with length-polymorphic (*msp-1*, *msp-2*, *glurp*) (3, 7–9) and microsatellite (4, 10) approaches.

Deep sequencing of highly SNP-polymorphic amplicons (AmpSeq), is an attractive next-generation methodology, experimentally shown to have much greater ability to detect low-density clones than existing methods based on length-polymorphic genotyping. This has the potential to substantially improve molecular correction over existing methods. AmpSeq methods are well established in the wider malarial context for tracking specific genes (e.g., for drug resistance) within populations (11–13), and for evaluating the efficacy of the RTS,S/AS01 vaccine (14). AmpSeq as a method to genotype malaria parasites in clinical trials is relatively novel (15) and, despite its putative advantages, has not yet been used as the primary genotyping endpoint for a clinical trial or TES (although it has been used to reanalyze archived blood samples (15)).

This paper uses an *in silico* pharmacological approach to evaluate the putative advantages AmpSeq for improving molecular correction. This builds on our previous work using mechanistic pharmacokinetic/pharmacodynamic (mPK/PD) modelling of malaria drug treatment to produce simulated trial data in which the ‘true’ underlying failure rate in the simulation is known. This allowed a critical appraisal of existing TES methodology (3, 4, 16) and enabled us to quantify the accuracy of molecular correction based on length-polymorphic and microsatellite markers. A reasonable hypothesis is that the more accurate and sensitive genotyping afforded by AmpSeq will lead to more accurate efficacy estimates in clinical trials and TES than existing, widely used methods mandated by the WHO and CDC. Here, we apply this mPK/PD modelling approach to quantify the accuracy of efficacy estimates produced using AmpSeq data of 5 SNP-polymorphic markers/loci. Our aim is to identify any potential problems and pitfalls *in silico* before they occur *in vivo*, optimise the likely use of AmpSeq for molecular correction in regulatory trials, and suggest appropriate guidelines for their deployment.

The main advantage of AmpSeq over existing methods is its increased sensitivity i.e. increased ability to detect genetic signals from malaria clones present at low densities in human blood samples. Sequencing may achieve substantial read depths but a practical problem in processing these reads is to distinguish low-number reads from artefacts. This requires a bioinformatics cut-off (BIC) value, below which low-number reads are regarded as artefacts and ignored. In the case of malaria it appears that BIC=1% is the most robust value, as extensively discussed elsewhere (17) although we also investigate a (hypothetical, perfect) BIC-> 0% and an alternative BIC of 2%. In all cases BIC should be read as the sensitivity of the method to detect low-density malaria clones e.g. with BIC=1% then any malaria clone(s) whose number/density is less than 1% of the total parasitaemia in a human will remain undetected.

## 2. Summary of Materials and Methods

### Simulating parasitaemia and their genotypes in therapeutic efficacy trials

Published mPK/PD methodologies (3, 4) were used to generate datasets of parasite numbers over time post-treatment for 5,000 simulated patients treated with either Dihydroartemisinin-Piperaquine (DHA-PPQ) or Artemether-Lumefantrine (AR-LF). These simulated patients differed in key factors related to their parasite dynamics post-treatment i.e. their individually-assigned PK parameters, the level of resistance in the patient’s parasites (the PD element), their parasitaemias at treatment, the local intensity of transmission and so on, as discussed later. Note that mPK/PD methodology tracks number, rather than density, of parasites; see Supplementary Material, Part 1 for how to interconvert these metrics. The simulations produce parasitaemia post-treatment for each patient simulated in the trial. Specifically, we track individual parasite clones present at treatment (which may be cleared or recrudesce) and of new infections.

These simulated patients were followed up for 42 days following DHA-PPQ treatment or 28 days following AR-LF treatment and ‘tested’ for recurrent parasitaemia by light microscopy on the scheduled days of follow-up i.e. days 3, 7 and every week thereafter in line with WHO guidelines (18). For each patient, our model calculated the day of follow-up on which recurrent parasites were first detectable (noting that some patients will never show recurrence) according to the procedure described in (3).

Genetic diversity of infections at treatment is termed the Multiplicity of Infection (MOI) which is the number of (detectable) genetically-distinct parasite clones in the blood sample. MOI depends on intensity of transmission so “high” and “low” MOIs were explored. Each parasite clone within the MOI had a total number of parasites drawn from a log-uniform distribution according to two ranges: 10^10^ to 10^11^ (the default range) and 10^8^ to 10^11^ (for sensitivity analysis). Genetic diversity in recurrences depends on the Force of Infection (FOI) which is the rate at which new infections become established in a patient. We incorporated FOI as the mean of a Poisson distribution from which the day(s) of new infections are randomly selected for each patient. The FOI means were 0, 2, 8 and 16 per year, broadly representing areas with no, low, medium and high ongoing transmission, respectively. New infections emerge from the liver as cohorts of 10^5^ total parasites and their fate (cleared or survive) depends on drug concentrations in that host on the days following their emergence from the liver (many emergences that appear shortly after treatment will be killed by the highly persistent partner drugs PPQ or LF). Each malaria clone at treatment or recurrence had genotypes defined at five AmpSeq markers (*cpmp*, *ama1-D3*, *cpp*, *csp* and *msp-7*) with allelic diversity obtained from (15). We use the first three markers by default but also included the less diverse markers *csp* and *msp-7* in our simulations as these may be used when other markers fail to amplify (15) and allows us to evaluate whether increasing the number of *Ampseq* markers would significantly improve accuracy.

A technical description of this methodology, including mPK/PD parameters and references to previous work is provided in the Supplementary Material, part 1. All simulations were conducted using the programming language R (version 1.2.5001) (19). The simulated data from the trial was then processed as follows.

### Allele detection in simulated datasets: the blood sampling limit and bioinformatics cut-off (BIC)

Our model determined which AmpSeq alleles were detected in initial and recurrent samples with a modified version of the methodology described in (3, 4) and described in the Supplementary Material, part1. It follows a two-stage process as follows:

1. A clone must have a sufficiently high parasitaemia that infected erythrocytes physically enter a finger-prick blood sample; this is obviously a prerequisite (but not a guarantee) for their later detection by PCR. This limiting level of parasitaemia is termed the ‘blood sampling limit’ (3, 4) and two limits were utilized: 10^8^ total parasites as used previously (3), and 10^7^ total parasites.
2. An amplicon allele will only be identified by the bio-informatics pipeline as present in the blood sample if it exceeds an empirically-determined threshold number and/or proportion of total reads (if below this threshold, the reads are regarded as “noise”). We call this threshold the bioinformatic cut-off (BIC) inherent in the experimental protocol. In our previous work [15], only reads present at >1% were regarded as “real” as this is highly robust [15]; consequently we use this value (i.e. BIC=1%) as the baseline for our simulations. We also investigated a more stringent threshold of 2% (i.e. BIC=2%). Since BIC is experimentally determined, we must anticipate settings where, for example, researchers are sufficiently confident in their technology and results that they regard reads present at proportions above one in 500 of the total reads, or even above 1 in 1000 (i.e. BIC=0.02% or BIC=0.1% respectively) as confirming the presence of that AmpSeq allele. Also, future technical advances may allow researchers to reduce this threshold even further e.g. to BIC=0.001. Rather than investigate all these BIC thresholds separately (e.g. BIC=0.2%, 0.1%, 0.001%, etc) we take the approach of investigating how the results would change as the threshold became extremely low i.e. BIC=0.0000….1 i.e. as BIC tended to zero (BIC->0). The logic is that if there is little improvement in accuracy as BIC falls from BIC=1% to BIC->0, then other values (such as the exemplar values BIC=0.2%, 0.1%, 0.001%) would also have little impact.

### Matching threshold for molecular correction and subsequent estimation of failure rate

Classification of recurrences as drug failures (i.e. recrudescences) or new infections was primarily based on the three most diverse markers *cpmp*, *cpp* and *ama1-D3*. A ‘matching threshold’ is used to classify a recurrence as a recrudescence when the number of markers with at least one shared allele between the initial and recurrent samples were greater than or equal to the matching threshold. For example, a threshold of ≥2 meant the initial and recurrent samples must share alleles at 2 or more markers to classify the recurrence as recrudescence. Recurrences below this threshold were classified as new infections. Once all recurrences are classified as recrudescences or new infections, the drug failure rate estimates in the trial were calculated using survival analysis, as per WHO procedure (18).

### The potential impact of gametocyte genotyping signals

Mature falciparum infections often contain relatively high densities (up to 10% of total parasitaemia) of their transmission stage, gametocytes. These stages do not cause symptoms, are unaffected by most drugs, and decline slowly post-treatment with a half-life of around 2 to 6 days. This slow decline has raised concerns that their genetic signals could be detected when genotyping recurrences using highly sensitive genotyping techniques such as AmpSeq. These signals would be mistaken for signals arising from persisting asexual stages, would be counted in the matching threshold described above, and hence bias molecular correction by increasing the likelihood of a ‘match’ between original and recurrent infections i.e. will potentially generate “false positive” recrudescences. Once we know the number of gametocytes at treatment, the lag time before they start to decline, and the rate of decline thereafter, it is straightforward to track their numbers following treatment and hence their potential detection and impact on molecular correction. However, discussions of calibration, algebra, and presentation of results became rather lengthy so, to maintain focus in the main text, we describe how we simulate their likely impact in a stand-alone Supplementary Material, Part 3.

We provide a summary of the parameters used in the simulations, and their values, in Table 1.

**Table 1.**
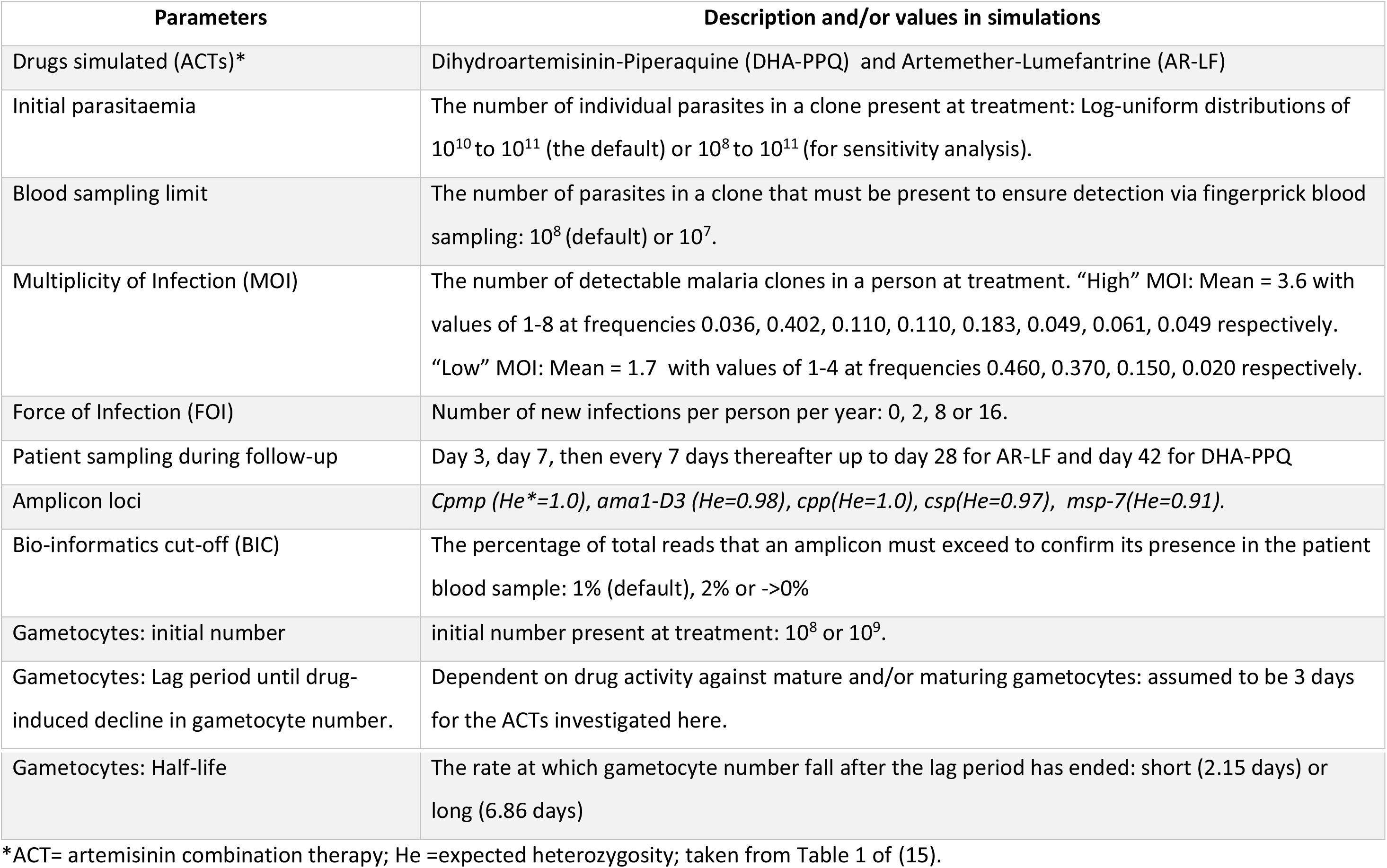
Summary of parameters used in the simulation and their values.

## 3. Results

Failure rate estimates for simulated populations with varied MOI and FOI treated with DHA-PPQ or AR-LF are shown in Figure 1. Three important model parameters were used as baseline scenario: BIC=1%, a “blood sampling limit” of 10^8^ total parasites, and initial parasite number drawn from a log-uniform distribution between 10^10^ and 10^11^. The true failure rate for DHA-PPQ was 11.6% and 6.3% in areas of high and low MOI respectively (true failure rates increase with MOI as the drug has to successfully clear more initial clones). The corresponding true failure rates for AR-LF were 12.2% and 8.2%.

**Figure 1.**
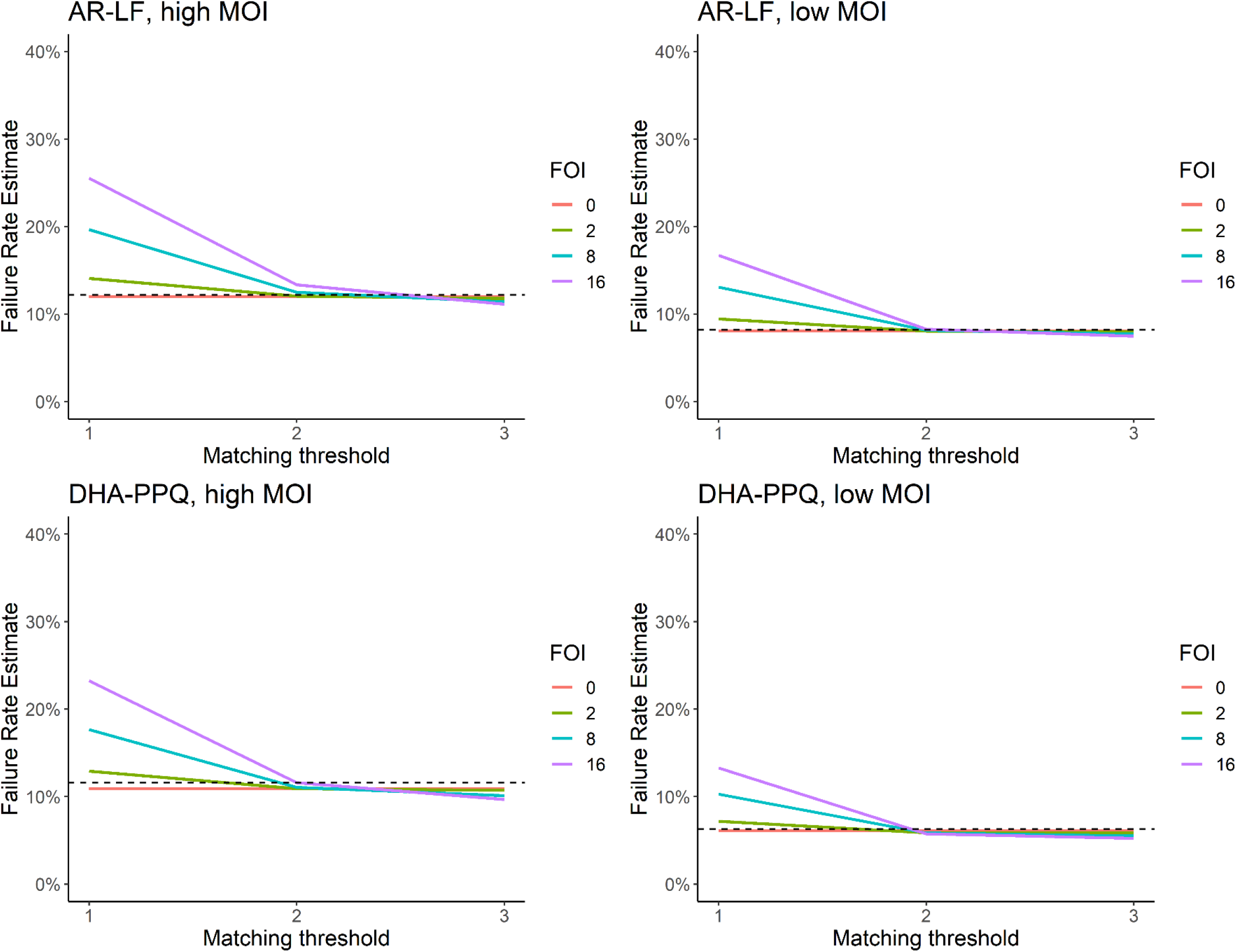
Failure rate estimates obtained using AmpSeq for Dihydroartemisinin-Piperaquine (DHA-PPQ) and Artemether-Lumefantrine (AR-LF) in setting of low and high Multiplicity of Infection (MOI) and a range of Force of Infection (FOI) values. Genotyping is based on 3 AmpSeq markers. The X axis shows the matching threshold used to define a drug failure (e.g. for a threshold of 2, then alleles at 2 or more loci must match in the initial and recurrent sample). The true drug failure rate is marked by the horizontal dashed black line.

A matching threshold of ≥2 or =3 AmpSeq loci used to classify recurrent infection as recrudescence produced highly accurate failure rate estimates (Figure 1) and performed much better than previously observed in our analysis of similar matching thresholds based on length-polymorphic WHO-recommended markers, and for microsatellites (3, 4). This was true under all scenarios i.e. for DHA-PPQ and AR-LF, in both high and low MOI and across all FOI values (Supplementary Material, Part 2). The accuracy of using three loci implies it is unnecessary to genotype more AmpSeq loci and this appears to be the case: genotyping 4 or 5 additional loci did not improve accuracy (Supplementary Material, Part 2).

An important operational question is whether technological advances capable of reducing BIC to below 1% will results in better estimates. This is unlikely given the accuracy of using BIC=1% (i.e. Figure 1) but we re-ran the analyses using the theoretical minimum value of BIC->0% and, as expected, found no improvement. Notably, increasing BIC to 2% also had a negligible impact on accuracy (Supplementary Material, Part2).

Sensitivity analyses were conducted by repeating simulations with altered model parameters to confirm their values did not affect the conclusions. Lower blood sampling limit or increasing initial parasite distributions showed no qualitative differences and negligible quantitative differences to results shown on Figure 1 (Supplementary Material, Part 2).

## 4. Discussion

Existing research has identified suitable SNP-polymorphic AmpSeq loci for genotyping malaria parasites (14, 17) and confirmed and quantified their superior ability to detect low-density clones compared to traditional length-polymorphic genotyping methods (15, 17, 20). AmpSeq also provided improved estimates of MOI (21) and identified appropriate thresholds for allele detection (17, 22). Here, we aimed to quantify the hypothesized increase in the accuracy of failure rate estimates in clinical trials (and eventually TES) that should result from AmpSeq’s increased ability to detect low density clones.

Our simulations show that genotyping 3 AmpSeq markers (*cpp*, *cpmp*, *ama1-D3*) using a matching threshold (number of markers with a shared allele between initial and recurrent samples) of ≥2 or =3 to classify recurrences as recrudesces returns highly robust estimates under a range of plausible MOI distributions and FOI values (Figure 1 and Supplementary Material, Part 2). Gruenberg et al. (15) noted that using a matching marker threshold of ≥2 is analogous to the new “≥2/3 threshold” being proposed for molecular correction with *msp-1/msp-2/glurp* length-polymorphic genotyping (3, 5) while a threshold of =3 is analogous to the 2007 WHO consensus methodology (2). It has been proposed that applying the ≥2/3 threshold to length-polymorphic markers leads to significantly increased accuracy of failure rate estimates (3, 7); the analogous ≥2/3 approach applied to AmpSeq returns even better, highly accurate failure rate estimates. The advantage of using AmpSeq over other markers is therefore due to their improved ability to detect genetic signals from low density clones.

The results presented here assume alleles can only be detected at frequencies >1% within a sample (i.e., BIC=1%). In reality, experimental mixtures suggest that AmpSeq is potentially even more sensitive than this but a BIC=1% is required to avoid inclusion of PCR errors / artefacts and environmental contaminations (15),. A value of BIC=1% reflects present technology for robust genotyping (15) but we wished to anticipate and evaluate technological advances that may reduce this limit. Our results show that reducing BIC to the hypothetical perfect detection limit as BIC->0% (and assuming no false positive occurred) *in silico* made negligible difference to the accuracy of the method; this most likely occurs because low-density clones that could be detected by the hypothetical perfect BIC->0% are likely to be below the blood sampling limit i.e. are unlikely to physically enter the finger-prick samples used in clinical trials and routine TES studies. The additional results (Supplementary Material, Part 2) also showed that BIC could be increased to 2% with negligible reductions in accuracy of molecular correction. Many surveys have shown a high prevalence of extremely low-density clones by deep-sequencing venous blood samples whose volumes are several magnitudes larger than a finger-prick. Whether resistant clones present at such extreme low-density are likely to recrudesce after treatment is unknown (one argument is that they are controlled by host immunity so are unlikely to recrudescence). Unfortunately, we cannot genotype such low density clones and test this directly in field trials using current technology because, as described above, the sequence depth is not the issue; any amplification step in the genotyping protocol will limit the sensitivity to around BIC=1% to avoid contamination producing too many low-density false positive haplotypes calls. Furthermore, setting BIC=1% reduces the risk of detecting genotypes of gametocytes persisting from treatment; the dangers of this have been raised previously (e.g. (15, 23)) and are quantified in our Supplementary Material, part 3.

The results emphasise that quality rather than quantity of amplicon loci is the key to obtaining accurate drug failure rates. Relatively large suites of amplicon markers (often 100+) are being developed for population genetic analyses of species including *Falciparum* spp but these require strict quality control before being used for molecular correction. This is why our previous work was so strict in this respect (15) i.e. taking three independent replicates per dried bloodspot sample and only classing an allele as present if (i) it was detected in all three replicates at more than 1% of total reads (i.e. BIC=1%) (ii) there were a minimum of 10 reads per allele and 50 reads per amplicon in each replicate. The final check before using amplicons for molecular correction is to confirm there is sufficient local genetic diversity (quantified as expected heterozygosity) at the amplicon to ensure that there is only a small probability of two independent blood samples sharing the alleles purely by chance. Finally, assuming high levels of genetic diversity, using more than around 5 amplicons in our simulation resulted in little, if any, improved accuracy (and may even increase the risk of genotyping errors).

Traditional molecular correction with length-polymorphic markers has been conducted using either gel-based or capillary electrophoresis (CE) of PCR products. The 2007 WHO guidelines contained protocols for both gel-based electrophoresis and CE (2, 9) but CE-based sizing using an automated sequencer offers higher sensitivity and ability to discriminate between alleles with minimal size differences; CE is now widely used and has generally phased out gel-based electrophoresis for molecular correction (9, 17). However, this improved discriminatory power is unable to compensate for the key deficiency of such methods, i.e. the inability to detect genetic signals from low-density clones, and our results clearly demonstrate the superiority of using amplicons for molecular correction. The adoption of AmpSeq will require use of next-generation sequencing platforms. The economic cost of reagents and deploying these machines to sub-Saharan Africa and South East Asia for use in malaria surveillance is likely to be significant and will necessarily need special training for sequencing data analysis with existing software e.g. HaplotypR, SeekDeep and PASEC (17, 22, 24), reagent supply, and equipment maintenance. Having AmpSeq facilities in every sentinel site is unlikely to be feasible, particularly in the short term. The future putative deployment of AmpSeq for analysis of routine malaria TES would probably require equipping a central site – one per country or even regionally if necessary, with the technology and expertise required to implement the methodology. Economic and technological factors appear to be the largest obstacle for AmpSeq deployment as a molecular correction methodology in routine TES surveillance but should be balanced against the long-term economic benefits of generating the accurate failure rate estimates required to ensure a sustainable supply of effective antimalarial drugs.

Finally, we stress that we have presented *in silico* simulations that we believe are a necessary pre-requisite for understanding how AmpSeq genotyping is likely to behave during their analysis in real clinical trials or TES studies for molecular correction. There is no guarantee that what occurs *in silico* will fully reflect what occurs in practice but our results strongly suggest that simple analysis of three to five genetically-diverse Ampseq loci should provide a baseline platform to obtain accurate drug failure rate estimates in trials and TES. This will need to be tested in early trialanalysis. For example, researchers may choose to genotype a larger bank of AmpSeq markers in which case our results suggest choosing the best (i.e. reliably genotyped, genetically-diverse loci) three to five loci should be sufficient to obtain good failure rate estimates. Similarly, genotyping technology may improve such that BIC may routinely fall below 1%. Analyses may improve by incorporating the probability that a recurrence can match the treatment genotype purely by chance (which is low for the genetically diverse Ampseq we simulate). Finally, designing a Bayesian analysis of AmpSeq genotyping (as previously developed for microsatellites in *P. falciparum* (10) and similar outcomes such as recurrences in *P. vivax* (25)) may be more accurate than simple counting the number of matches. It remains to be seen whether these improvements will have a significant, or a negligible, impact on failure rate estimates. One danger of more sophisticated analyses is that researchers often do not engage with the processes: they either ignore them in favour of simpler methods, or the implementation is so complicated that misunderstanding arise. For example, a recent review (26) showed that many TES performed in sub-Saharan Africa did not conform to current WHO guidelines for classifying recurrences as drug failures or new infections based on simple identification of shared alleles. Subsequent analysis frequently regarded new infections as equivalent to drug cure (26), again contrary to WHO guidelines that recommend their removal from the analysis or incorporation into survival analysis. It is important to recognise that in-country researchers generating genotyping data naturally wish to analyse their own data, rather than passing it to external collaborators, which is why simple counting is operationally preferable to the slightly more accurate Bayesian analyses that are likely to be technically inaccessible to most groups.

In summary, the results presented here suggest AmpSeq is a strong candidate to become the new gold standard method of molecular correction in malaria clinical trials as such studies would be supported by the requisite technical expertise. As infrastructure and technical expertise increases in endemic countries, the use of amplicons may be eventually become feasible in routine TES. Simulations suggest that accuracy would not improve significantly from the current methodology even if a hypothetical perfect detection of low density clones was achievable (i.e. BIC->0%) implying that AmpSeq is unlikely to be usurped by newer, more sensitive genotyping technologies. In fact, further increasing sensitivity (i.e. reducing BIC) may potentially become counter-productive through capturing false positive signals from gametocytes (Supplementary Material, Part 3). Analysis of AmpSeq data is also straightforward and can use simple counting of allelic matches to distinguish recrudesces from new infection.

## Supporting information

Combined Supplemental Material

## Conflict of interest statement

None of the authors have any associations that might pose a conflict of interest with authorship of this manuscript

## Funding

This work was supported by the UK Medical Research Council (grant numbers G1100522 and MR/ L022508/1); the Bill and Melinda Gates Foundation (grant number1032350); the Malaria Modeling Consortium (grant number UWSC9757).

## Notes

### Competing Interest Statement

The authors have declared no competing interest.

