## Supplementary material for "Should deep-sequenced amplicons become the new gold-standard for analysing malaria drug clinical trials?": Combined Supplemental Material

### Combined Supplementary Material

This document contains three independent SI documents (identified by their footers) i.e. with their own tables, figures, citations.

Note that SI part 1 cross-references figures in SI part 2.

#### Supplementary Material #1: Terminology and technical details of the pharmacological simulations.

There is potential confusion in the use of identical terms to mean different things in the sequencing and "classical" genetic fields. We therefore clarify the terminology used in this paper.

In classical genetics a haplotype can refer to the full genetic complement of a (haploid) malaria parasites, as can the term "genotype". Some existing AmpSeq literature refers to individual gene variants as "haplotypes" or microhaplotypes (see, for example, (1-3)) to distinguish them from "genotypes" which is used to refer to a SNP. Similarly, Gruenberg et al. (4), (which we reference heavily because we use their methodology as the basis for our *in silico* analysis methods) defined a haplotype as "a unique sequence variant of an entire amplicon". Elsewhere, AmpSeq literature may use the term haplotype in the more traditional sense i.e. to describe a group of alleles that are inherited together (5).

In this manuscript we deliberately avoid the ambiguous term "haplotype" and use "alleles" to refer to individual gene variants. This avoids potential confusion, is consistent with "classic" genetic terminology, and ensures consistency of terminology with previous investigations of different markers used for molecular correction (i.e., length-polymorphic markers and microsatellite markers (6, 7)).

Mechanistic Pharmacokinetic / Pharmacodynamic (mPK/PD) models use PK/PD equations to quantify drug concentration and drug killing and create models of parasite dynamics (numbers) over time post-treatment. These differ from "traditional" PK/PD modelling which analyses *in vivo* data of drug concentration and their effects using sophisticated modelling methods to elucidate underlying PK/PD parameters. Such parameters can subsequently be used to calibrate mPK/PD models, as we do here.

##### 1.1 Pharmacokinetics: partner drug and Artemisinin concentrations over time.

Five thousand patients were dosed with DHA-PPQ or AR-LF using the PK parameters shown in Table S1.1. Drug dosing for DHA-PPQ and AR-LF (Table S1.2) reflect the recommended dosing regimen published by the WHO in 2015 (8). in the mPK/PD models used here each patient is given a precise dose according to their body weight (noting that, in practice, tablets contain a fixed weight of drugs so doses are given according to patient age or weight bands.

The drug concentration over time profiles produced by the model for PPQ (assuming a two-compartment model for PPQ) and LF (assuming a one compartment model for LF) are given in Figures S1.1 and S1.2. The equivalent profiles for DHA and AR are given in Figures S1.3 and S1.4. The methodology used to generate these concentrations is described in (9).

**1.2 Pharmacodynamics: and relationship between drug concentration and parasite killing.**

The PD parameters determine the rate of parasite killing (at a given concentration of drug) for each parasite clone; the mechanistic relationship between these parameters and drug killing is described in (9-11) (specifically in (9)). The mPK/PD method requires three parameters: The maximal parasite killing constant, Vmax, the slope factor, *n*, and the "half maximal inhibitory concentration" (IC50), which is the concentration of drug at which half-maximal parasite killing occurs. These parameters are given in in Table S1.3.

To calibrate failing drugs, the IC50 of PPQ could be obtained from *in vivo* data, but the IC50 of "failing" LF is a hypothetical value determined by us that results in a ~10% failure rate in the mechanistic model. In previous work using mPK/PD models we simulated both failing and non-failing drug calibrations (6, 12); in this paper we only simulate failing drugs because we are interested only in molecular correction, not other factors such as appropriate duration of follow-up. We have also previously altered drug IC50 to allow for  changing MOI (see below); higher MOI (all other things being equal) will lead to higher failure rate as there are more initial infections to be cleared by treatment, so IC50 was previously increased in lower MOI settings to keep failure rates ~10% (6). Such an approach has led to confusion in the past, so here we opt to use a single IC50 value for a partner drug with the understanding that there will be a slightly higher true failure rate in higher MOI scenarios.

**1.3 Multiplicity of infection (MOI).**

Multiplicity of infection (MOI) is the number of genetically distinct malaria clones in a patient's blood sample.  Unsurprisingly, MOI increases with malaria transmission intensity so two MOI distributions were modelled. A "high MOI" was representative of the MOI in an area of intense transmission, in this case Tanzania where MOIs of 1-8 were assigned with probabilities 0.036, 0.402, 0.110, 0.110, 0.183, 0.049, 0.061, 0.049 respectively (12). A "low MOI" distribution was based on data from Papua New Guinea with probabilities of 0.460, 0.370, 0.150 and 0.020 for an MOI of 1-4 respectively. These two distributions were subsequently used to check if the accuracy of molecular correction was consistent between high/low MOI.

These MOI distributions are identical to those previously used for simulation of length-polymorphic markers *msp-1*, *msp-2* and *glurp* (6). Note that the MOI distributions were derived, in the first instance, from in vivo data using length-polymorphic markers, not AmpSeq. Given that AmpSeq provides a higher detectability of minority clones (2, 13), MOI estimates from a given population with AmpSeq should be higher than MOI estimates with length-polymorphic markers. However, use of AmpSeq for this purpose is extremely novel, and useable MOI distributions obtained with AmpSeq are limited. To the best of our knowledge, the only available MOI distributions obtained using AmpSeq are across multiple countries or study sites (5) and would not be appropriate to generate MOI distributions.

**1.4 Force of infection (FOI).**

The force of infection (FOI) is the rate at which patients in endemic countries acquire new malaria infections. We incorporate FOI by defining the mean of a Poisson distribution from which the number of reinfections that occur (per year) in a patient is randomly selected. The number of reinfections is then scaled to reflect the time-frame of the follow-up period (i.e., an FOI of 12 in a year would relate to an average of 1 reinfection in a 4 week follow-up period). The FOI parameters investigated were 0, 2, 8 and 16, broadly representing an area with no, low, medium and high ongoing transmission, respectively. A full discussion of the relationship between FOI and important epidemiological parameters such as the annual entomological inoculation rate (aEIR) can be found in the supplementary material of (6).

##### 1.5 Initial parasite numbers and relationship with parasites densities.

We quantify parasitaemia as the total number of parasite in the patient, rather than parasite densities in blood samples (as previously e.g.(12)). We do not parameterize patients in a way that would allow us to easily convert total numbers to parasite densities (i.e., patients do not have parameters for blood volume, white blood cell [WBC] count, red blood cell count, etc.), nor would including these parameters aid the mechanistic simulation of the model or improve the accuracy of the results. For reference, assuming a patient with 4.5 litres of blood and a WBC count of 8,000/µl of blood, parasitaemias of $10^{10}$ and $10^{11}$ would correspond to densities of 2,222 parasites/µl of blood and 22,222 parasites/µl of blood, respectively, according to the WHO counting procedure (14). Previous modelling approaches used $10^{12}$ parasites as the upper limit of parasitaemia; this level of parasitaemia is likely to be lethal or at least exceed the maximum parasite density exclusion criterion in a clinical trial (typically 100,000 parasites/µl); hence, we used $10^{11}$ as the upper limit for any single clone at the time of treatment. A value of $10^{10}$ was used as the lower limit in previous work on length-polymorphic and microsatellite markers (6, 12) in order to correctly represent the MOI because although it is possible for patients to harbour low-density clones, these clones would not be detected with these methods and consequently not be included in the MOI count. AmpSeq improves detection of low density clones so, to avoid any doubt, we model a lower limit of $10^8$.

##### 1.6 Detection of recurrence during follow-up.

The model checked each day of scheduled follow-up to determine whether a patient had parasitaemia sufficiently high that a recurrence would be detectable by light microscopy (LM). Detection by LM was assume to occur if the total number of parasites in the patient was $\geq 10^8$ on that day (15). This corresponded to a parasite density of roughly 20 parasites/µl of blood. Detection by LM varies according to the skill of the microscopist (14) and this this limit reflects that of an “expert” microscopist

If total parasitaemia exceeded $10^8$ on day 3 but was <25% of the total parasitaemia of the initial sample, the patient continued in the trial; if parasites were present at >25% of initial parasitaemia, that patient was classed as an early treatment failure and withdrawn from the study as per WHO procedure (16). Note that for subsequent calculations and analysis, an early treatment failure is considered a recrudescence.

##### 1.7 The blood sampling limit

A finite volume of blood enters the genotyping procedure. A parasite clone will remain undetected if its density is so low that none of its constituent parasites were physically included in the blood sample analysed. Thus, the parasite density and volume of the processed blood sample defined the limit of detection and we quantify this as the “blood sampling limit”. Obviously, this blood sampling limit differs between methods and laboratories. Typically, the equivalent of 1 µl of whole blood is introduced into a PCR. Assuming 5 L of blood in the human body gives a total of $5 \times 10^6$ µl of blood. For a clone to be detected a minimum of 1 parasite (which carries a single DNA template) would need to be present in 1 µl of blood so there would need to be at least $5 \times 10^6$ of a given clone present for that clone to be physically sampled in the genotyping process. In practice, we need to allow for the fact that sub-optimal storage conditions (such as temperature) frequently occurs in the field, which can lead to DNA template breakages. Finally, there is periodical absence of sequestered parasites from the peripheral blood. Consequently, the limit of detection will be much higher than 1 parasite per 1µl of blood. It was therefore assumed 10 to 20 parasites per µl would be required to reliably ensure its detection, corresponding to a total parasitaemia of 5 x $10^7$ to $10^8$; the upper limit i.e. $10^8$ was selected to ensure reliable detection of that clone and because it is consistent with the microscopy detection limit.

The blood sampling limit was lowered to $10^7$ as part of a sensitivity analysis on model parameters.

##### 1.8 Genetic diversity at Ampseq loci.

Each malaria clone in patients in our simulated data-sets had genotypes defined by alleles at 5 AmpSeq markers (*cpmp*, *ama1-D3*, *cpp*, *csp* and *msp-7*). Genetic diversity at these five markers was obtained by data presented in (4) which included (i) Details of the identification, sequencing and additional information for the 5 markers *ama1-D3* (PF3D7_1133400), *cpmp* (PF3D7_0104100), *csp* (PF3D7_0304600), *cpp* (PF3D7_1475800) and *msp-7* (PF3D7_1335100) (ii) A summary of each marker, including expected heterozygosity (He) as given in their Table 1 (iii) full allele frequency distributions which are also included in their supplemental material. Note that the diversity of each marker *in vivo* will vary between TES sites, but diversity can be assessed bioinformatically for each study site from the global *P. falciparum* genome in the MalariaGen data base i.e. https://www.malariagen.net/data/terms-use/p-falciparum-community-project-terms-use

#### Part 2: Additional Results and discussion

##### 2.1 Additional results obtained when changing model parameters

Four important model parameters were varied to assess their impact on results: The BIC value, the blood sampling limit, the lower limit of the parasite number distribution in the patient sample taken at treatment, the number of *Ampseq* loci genotyped.

##### 2.1.1 Changing the bioinformatics cut-off (BIC) value.

(see Figures S2.1 and S2.2)

The value BIC=1% was used in the baseline calculations (i.e., minority alleles were detected if they exceeded 1% of total reads). This threshold is user-defined by the genotyping software when AmpSeq markers are genotyped (3) so assessing the impact of this parameter on the accuracy of molecular correction is important. As noted by Gruenberg et al., “a stringent cut-off is required for excluding sequencing errors; on the other hand, a less stringent cut-off would be desirable for maximized detection of minority clones”. Here, we increased this threshold to 2% to assess the impact of a higher cut-off point. We also reduced it to 0% to assess the difference between 1% and 2% cut-offs from the hypothetical perfect detection of minority clones that would occur at BIC->0%, noting that 0% is unfeasible in practice because such a cut-off permits the inclusion of sequencing errors and contaminations.

Failure rate estimates for DHA-PPQ and AR-LF with low and high MOI and a range of FOI values are shown in Figure S2.1 for BIC->0% and Figure S2.2 for BIC=2%. When compared to BIC=1% (Figure 1 of main text), a BIC->0% resulted in slightly higher failure rate estimates and BIC=2% resulted in slightly lower failure rate estimates. In both cases, the difference was negligible and failure rate estimates obtained using BICs of ->0%, 1% and 2% were very close. In other words, the currently proposed threshold of BIC=1% has negligible difference to a hypothetical perfect detection scenario. If genotyping errors were to necessitate a higher BIC, our results indicate that BIC=2% could be used with minimal loss of accuracy compared to BIC=1%.

###### 2.1.2 Changing the blood sampling limit

(see Figures S2.3)

The blood sampling limit (the parasitaemia of a clone required for it to be physically included in a finger-prick blood sample used for genotyping) was $10^8$ total parasites in the baseline model. This limit was calculated based on realistic blood sampling processes (see Supplementary Material, Part 2) but given the ability of AmpSeq to detect low frequency alleles, it was necessary to check this assumption was not biasing results. The blood sampling limit was therefore reducing to $10^7$ total parasites (i.e. a lower density clones would be included in the blood sample).

Failure rate estimates obtained with the lower blood sampling limit are shown for DHA-PPQ and AR-LF in Figure S2.3. Results were qualitatively extremely similar to the baseline model (Figure 1 of main text). There was an extremely small increase in failure rate estimates at higher FOI (8 and 16) of ~0.02% when using lower number of matches (≥1 or ≥2) to classify a recrudescence. In short, results were functionally identical to the baseline assumption of a blood sampling limit of $10^8$, so the assumed value of the blood sampling limit did not appear to affect the failure rate estimates obtained using AmpSeq markers.

###### 2.1.3 Changing the lower limit of the initial parasite number distribution

(see Figures S2.4)

The lower limit of the log-uniform distribution was changed from $10^{10}$ parasites (the baseline lower limit) to $10^8$ parasites (the upper limit remains as $10^{11}$ parasites). This allows us to investigate whether the accuracy of failure rate estimates generated using AmpSeq markers was affected by assuming a wider range of initial parasitaemia across clones (which will lead to increase proportions of low density clones). The true failure rate changed as the range of this distribution changed: 8.8% and 4.4% for DHA-PPQ in high and low MOI settings respectively and 10% and 6.6% for AR-LF in high and low MOI settings respectively. These true failure rates are slightly lower than the baseline scenarios, presumably because MOI is held constant so more low-density clones are present in the initial infections, and such low-density clones are less likely to recrudesce. However, the absolute change in true failure rate was negligible, so this effect did not appear to be large. The BIC was 1% and blood sampling limit was $10^8$, both as for the baseline model.

Failure rate estimates using a wider initial parasite number distribution (i.e. reducing log distribution of initial parasite number down to $10^7$) are shown for DHA-PPQ and AR-LF in Figure S2.4. Failure rate estimates were slightly lower in both cases, though this should be considered relative to the slightly lower true failure rate. The difference between each estimate and the true failure rate, and thus the qualitative conclusions, were identical to the baseline model i.e. classifying a recrudescence at ≥2 or 3 matches accurately recovered the true failure rate for both drugs, both MOI settings and all FOI values.

###### 2.1.4 Increasing the number of Ampseq loci

(see Figures S2.5 and S2.6)

We calculated failure rate estimates using 4 AmpSeq markers (including *csp*) and 5 AmpSeq markers (including *csp* and *msp-7*) – this represents inclusion of the less diverse markers in the marker data set. Failure rate estimates obtained using 4 or 5 AmpSeq markers are shown in Figures S2.5 and S2.6 respectively. In both cases, classifying a recrudescence at ≥2 matches no longer produces accurate failure rate estimates (compared to using 3 AmpSeq markers (*cpp*, *cpmp*, *ama1-D3*)) and leads to over-estimation of failure rate at higher FOI. This effect arises because inclusion of less diverse markers increases the probability that reinfections share, by chance, a given number of alleles with clones from the initial infection. There is consequently a slightly increased misclassification of reinfections as recrudescences. Our results indicated that if genotyping were to be done on 4 AmpSeq markers, a recrudescence should be defined as ≥3 matches, and for 5 AmpSeq markers a recrudescence should be defined as ≥4 matches. In summary, genotyping 4 or 5 AmpSeq markers does not significantly increase molecular correction accuracy compared to genotyping using 3 AmpSeq markers. We do note the practical point that genotyping additional markers will be useful in the case that any other markers fail to amplify or are otherwise corrupted in the genotyping process. This suggests that the best threshold is ≥(n-1) where n is the number of markers genotyped. We also note that Bayesian analyses will increase discrimination between clonal genotypes (e.g. (7, 17)) and we await such analyses with interested. Ours results imply that simple counting returns such accurate results that remains to be seen whether the improved results that should be obtained by Bayesian analyses will make any difference to failure rates estimates obtained by simple counting.

The AmpSeq markers used in these simulations were selected in highly SNP-polymorphic regions so there is a high number of alleles for each marker. A lower allele diversity has only been observed for *csp* in one geographic area i.e. PNG (with only 3 alleles (3)). Populations with low genetic diversity have not been genotyped using these markers to date. If/when lower diversity data-sets become available, this modelling work can be repeated to quantify the accuracy of failure rate estimates in such areas. The impact of lower genetic diversity would be to increase failure rate estimates due to “identity by chance” i.e. more reinfections will be misclassified as recrudescence due to them sharing alleles purely by chance. In such areas, it may be necessary to consider a) genotyping a larger number of markers and b) to use higher (more stringent) thresholds of matching loci to classify a recurrence as a recrudescence (i.e., the ≥2/3 threshold may over-estimate true failure rate and a =3/3 threshold may be superior). (c) Develop Bayesian methods of analysis and test whether they provide a clinically-significantly improved failure rate estimate Use of AmpSeq in TES requires genotyping the initial blood samples and quantifying the level of genetic diversity of those samples, such that informed decisions around the total number of markers to analyse and the threshold chosen can be made. Obtaining accurate MOI and allele diversity estimates is possible with AmpSeq due to their high resolution (2). Estimates of FOI (this parameter can be derived from annual entomological inoculation rate (aEIR), see SI of (6)) should also be obtained where possible. This epidemiological information can then be used to optimize AmpSeq analysis for a given TES.

#### 2.2 Additional discussion of results.

##### 2.2.1 Impact of Multiplicity of Infection (MOI) and drug type.

The multiplicity of infection at time of treatment and drug used (DHA-PPQ or AR-LF) appeared to have little impact on the qualitative results (Figure 1 of main text and Figures S2.1 to S2.6 below); this is consistent with results obtained when simulating other types of marker (6, 12).

##### 2.2.2 Impact of Force of Infection (FOI)

The effect of the Force of Infection (FOI) is shown on Figure 1 of main text and Figures S2.1 to S2.6 below.

When FOI is zero, there is no possibility of re-infection and failure rate estimates did not change as the matching threshold was altered (an obvious result because, when FOI=0, all recurrences must be recrudescence). In these circumstances, a slight under-estimate of true failure rate (<1%) occurred in two scenarios:

(i) If the recrudescence never reaches patency during follow-up i.e. never grows to $>10^8$ total parasites which is the limit of detection by light microscopy, see methodology in (6). This means the patient is erroneously classed as having cleared their initial infection.
(ii) If the recrudescent clone was comparatively low-density clone at treatment so was not detected in the initial sample (thus the recurrence is misclassified as a reinfection).

The under-estimate was larger for DHA-PPQ scenarios than AR-LF – consistent with previous work indicating that some recrudescences occur later than 42 days after treatment with DHA-PPQ, but nearly all have occurred by 28 days following treatment with AR-LF.

As FOI increased, it started to have an impact because introducing reinfections had two consequences:

(i) There was a higher likelihood of a recurrence containing both recrudescent clone(s) and new infections, and thus a higher chance for recrudescent clones to be below the detection threshold in the recurrent sample (i.e., some truly recrudescent alleles may not be observed in the recurrence).

(ii) More clones, and hence alleles, are likely to be present in the recurrent blood sample so the chance of finding a match with the initial infection purely by chance is increased. This would result in a reinfection being misclassified as a recrudescence (noting this chance will be close to 0 when a match is required at all markers). This occurred at all matching thresholds, but misclassification of reinfection with lower thresholds resulted in higher failure rate estimates and is why failure rate estimates are highest with FOI 16 using a matching threshold of ≥1 but lowest with a threshold of =3.

This affect the relative performance of different matching thresholds as described in the next section.

##### 2.2.3 Choice of matching threshold to define a recrudescence

Figure 1 of main text shows that the impact of FOI is heterogeneous at the different matching thresholds used to classify a recrudescence, so the key operational question is thus: What matching threshold produced failure rate estimate closest to the true failure rate, and is this threshold robust for multiple drugs and in multiple MOI and FOI settings? The results can be summarised as follows:

- A matching threshold of ≥1 appeared unsuitable because failure rates were severely overestimated at moderate or high FOI values (because there is high chance of a reinfection sharing, by chance, an allele present in the original infection, leading to reinfections being misclassified as recrudescences).
- A matching threshold of ≥2 greatly reduced the probability of matching-by-chance and returned accurate failure rate estimates.
- A matching threshold of =3 generally slightly under-estimated failure rates.

In summary, aside from the high MOI, FOI=16, AR-LF scenario (where a threshold of ≥2 caused a slight over-estimate of true failure rate), using a threshold of ≥2 appeared to be highly robust and returned the most accurate failure rate estimates in all scenarios. This is supported by results obtained when genotyping a larger number of Ampseq (see 2.1.4 above) where results indicated the most robust matching threshold was ≥(n-1) where n is the number of markers genotyped.

**References.**


1. Tessema SK, Hathaway NJ, Teyssier NB, Murphy M, Chen A, Aydemir O, Duarte EM, Simone W, Colborn J, Saute F, Crawford E, Aide P, Bailey JA, Greenhouse B. 2020. Sensitive, Highly Multiplexed Sequencing of Microhaplotypes From the *Plasmodium falciparum* Heterozygome. J Infect Dis corrected proof availble online.
2. Lerch A, Koepfli C, Hofmann NE, Kattenberg JH, Rosanas-Urgell A, Betuela I, Mueller I, Felger I. 2019. Longitudinal tracking and quantification of individual *Plasmodium falciparum* clones in complex infections. Sci Rep 9:3333.
3. Lerch A, Koepfli C, Hofmann NE, Messerli C, Wilcox S, Kattenberg JH, Betuela I, O'Connor L, Mueller I, Felger I. 2017. Development of amplicon deep sequencing markers and data analysis pipeline for genotyping multi-clonal malaria infections. BMC Genom 18:864.
4. Gruenberg M, Lerch A, Beck H-P, Felger I. 2019. Amplicon deep sequencing improves Plasmodium falciparum genotyping in clinical trials of antimalarial drugs. Sci Rep 9:17790.
5. Neafsey DE, Juraska M, Bedford T, Benkeser D, Valim C, Griggs A, Lievens M, Abdulla S, Adjei S, Agbenyega T, Agnandji ST, Aide P, Anderson S, Ansong D, Aponte JJ, Asante KP, Bejon P, Birkett AJ, Bruls M, Connolly KM, D'Alessandro U, Dobano C, Gesase S, Greenwood B, Grimsby J, Tinto H, Hamel MJ, Hoffman I, Kamthunzi P, Kariuki S, Kremsner PG, Leach A, Lell B, Lennon NJ, Lusingu J, Marsh K, Martinson F, Molel JT, Moss EL, Njuguna P, Ockenhouse CF, Ogutu BR, Otieno W, Otieno L, Otieno K, Owusu-Agyei S, Park DJ, Pelle K, Robbins D, Russ C, et al. 2015. Genetic Diversity and Protective Efficacy of the RTS,S/AS01 Malaria Vaccine. N Engl J Med 373:2025-2037.
6. Jones S, Kay K, Hodel EM, Chy S, Mbituyumuremyi A, Uwimana A, Menard D, Felger I, Hastings IM. 2019. Improving methods for analysing anti-malarial drug efficacy trials: molecular correction based on length-polymorphic markers *msp-1*, *msp-2* and *glurp*. Antimicrob Agents Chemother 63:e00590-19.
7. Plucinski MM, Morton L, Bushman M, Dimbu PR, Udhayakumar V. 2015. Robust algorithm for systematic classification of malaria late treatment failures as recrudescence or reinfection using microsatellite genotyping. Antimicrob Agents Chemother 59:6096-100.
8. World Health Organization. 2015. Guidelines for the treatment of malaria.
9. Kay K, Hastings IM. 2013. Improving pharmacokinetic-pharmacodynamic modeling to investigate anti-infective chemotherapy with application to the current generation of antimalarial drugs. PLoS Comput Biol 9:e1003151.
10. Hodel E, Kay K, Hayes D, Terlouw D, Hastings I. 2014. Optimizing the programmatic deployment of the anti-malarials artemether-lumefantrine and dihydroartemisinin-piperaquine using pharmacological modelling. Malar J 13:138.
11. Winter K, Hastings IM. 2011. Development, evaluation and application of an *in silico* model for antimalarial drug treatment and failure. Antimicrob Agents Chemother 55:3380-3392.
12. Jones S, Plucinski M, Kay K, Hodel EM, Hastings IM. 2020. A Computer Modelling Approach To Evaluate the Accuracy of Microsatellite Markers for Classification of Recurrent Infections during Routine Monitoring of Antimalarial Drug Efficacy. Antimicrob Agents Chemother 64:e01517-19.

13. Early AM, Daniels RF, Farrell TM, Grimsby J, Volkman SK, Wirth DF, MacInnis BL, Neafsey DE. 2019. Detection of low-density *Plasmodium falciparum* infections using amplicon deep sequencing. Malar J 18:219.
14. World Health Organisation. 2016. Malaria microscopy quality assurance manual – Version 2. World Health Organisation, Geneva.
15. Siahaan L. 2018. Laboratory diagnostics of malaria. IOP Conference Series: Environ Earth Sci 125:012090.
16. World Health Organization. 2009. Methods for surveillance of antimalarial drug efficacy. World Health Organisation, Geneva.
17. Taylor AR, Watson JA, Chu CS, Puaprasert K, Duanguppama J, Day NPJ, Nosten F, Neafsey DE, Buckee CO, Imwong M, White NJ. 2019. Resolving the cause of recurrent *Plasmodium vivax* malaria probabilistically. Nat Commun 10.
18. Kay K, Hodel EM, Hastings IM. 2015. Altering antimalarial drug regimens may dramatically enhance and restore drug effectiveness. Antimicrob Agents Chemother 59:6419-6427.
19. Staehli Hodel E, Guidi M, Zanolari B, Mercier T, Duong S, Kabanywanyi A, Ariey F, Buclin T, Beck H-P, Decosterd L, Olliaro P, Genton B, Csajka C. 2013. Population pharmacokinetics of mefloquine, piperaquine and artemether-lumefantrine in Cambodian and Tanzanian malaria patients. Malar Jl 12:235.
20. Saunders DL, Vanachayangkul P, Lon C. 2014. Dihydroartemisinin–Piperaquine Failure in Cambodia. N Engl J Med 371:484-485.

**Table S1.1** PK Parameter summary. A summary of the PK parameters used to simulate parasite dynamics post-treatment, adapted from Hodel et al. (10). The table shows mean values with coefficient of variation in brackets while square brackets are citations in support of the parameter values.

| **Drug** | **Dihydroarteminisin-Piperaquine (2 compartment model)** | | **Artemether-Lumefantrine** | | |
|---|---|---|---|---|---|
| | **DHA** | **PPQ** | **AR** | **DHA** | **LF** |
| **Vd (L/kg)** | 1.49 (0.48)(9, 10) | 346 (0.93)(18) | 46.6(0.82)(10) | 15(0.48) (9, 10) | 21(2.63)(9, 10) |
| **$Vd_1$ (L/kg)** | - | 443 (1.70)(18) | - | - | - |
| **ka (/day)** | - | 11.2 (2.17)(18) | 23.98(0.68) (9, 10) | - | - |
| **z (/day)** | - | - | 11.97(0.65)(9, 10) | - | - |
| **$Q_1$(L/day/kg)** | - | 69.7(1.01)(18) | - | - | - |
| **k (/day)** | 19.8(0.23)(10, 11) | 0.02*(18, 19) | - | 44.15(0.23) (9, 10) | 0.16(0.05) (9, 10) |

*PK: Pharmacokinetic, BW: Patient bodyweight, DHA: Dihydroartemisinin, PPQ: Piperaquine, AR: Artemether, LF: Lumefantrine, Vd: Volume of Distribution (central compartment for PPQ), $Vd_1$: Volume of Distribution (peripheral compartment), $Q_1$: Intercompartmental clearance (central-peripheral 1), ka: Absorption rate constant, z: Conversion rate of AR/AS into DHA, - : No data / not applicable.*

* *elimination rate for PPQ is calculated from clearance (CL) / Vd. CL is not shown here but is 4.5 * $BW^{0.75}$ as in (19); This means that elimination rate varies with body weight ( a common PK observation) so the value presented here is illustrative and represents a bodyweight of 42kg (the median bodyweight in previous studies* (18, 19)*). Partner drug IC50 values are not shown here; they vary between and within chapters, see individual chapters for these values. Piperaquine (PPQ) here follows a two-compartment model as described in Kay, Hodel & Hastings (18). Patient bodyweight (BW) in all simulations was drawn from a uniform distribution between 45-75 kg and is involved in the calculations for PPQ parameters (see* (18, 19)*).*

**Table S1.2** Drug dosing of the artemisinin and partner drug components of the ACTs for the mechanistic simulation of DHA-PPQ, AR-LF and AS-MQ

| **Drug** | **DHA-PPQ** | | **AR-LF** | |
|---|---|---|---|---|
| | **DHA** | **PPQ** | **AR** | **LF** |
| **Dose at 0 days (**mg/kg) | 4 | 18 | 1.7 | 12 |
| **Dose at 0.5 days (**mg/kg) | | | 1.7 | 12 |
| **Dose at 1 days (**mg/kg) | 4 | 18 | 1.7 | 12 |
| **Dose at 1.5 days (**mg/kg) | | | 1.7 | 12 |
| **Dose at 2 days (**mg/kg) | 4 | 18 | 1.7 | 12 |
| **Dose at 2.5 days (**mg/kg) | | | 1.7 | 12 |

*DHA: Di-hydroartemisinin, PPQ: Piperaquine, AR: Artemether, LF: Lumefantrine. Dosages listed are mg/kg, e.g., for a 45kg patient, a dose of 180mg of DHA would be given at each interval.*

**Table S1.3** A summary of the PD parameters used to generate parasite dynamics *in vivo* with an mPK/PD model. The table shows mean values with coefficient of variation in brackets while square brackets are citations in support of the parameter values.

| Drug parameter | Di-hydroarteminisin-Piperaquine (2 compartment model) | | Artemether-Lumefantrine | | |
|---|---|---|---|---|---|
| | **DHA** | **PPQ** | **AR** | **DHA** | **LF** |
| **IC50 (mg/L)** | 0.009 (1.17)(9, 10) | 0.02 (0.3) (20) | 0.0023(0.79)(9, 10) | 0.009(1.17)(9, 10) | 10 (1.02) |
| **Vmax** | 27.6 (9, 10) | 3.45 (11) | 27.6 (9, 10) | 27.6 (9, 10) | 3.45 (9, 10) |
| **n** | 4 (9-11) | 6 (11) | 4 (9-11) | 4 (9-11) | 4 (9-11) |

*IC50: Half maximal inhibitory concentration, Vmax: Maximal parasite killing constant, n: Slope factor. Half maximal inhibitory concentration (IC50) is shown for all artemisinins but not for partner drugs The coefficient of variation (CV) is provided in brackets where appropriate. Citations are provided in square brackets in support of parameter values.*

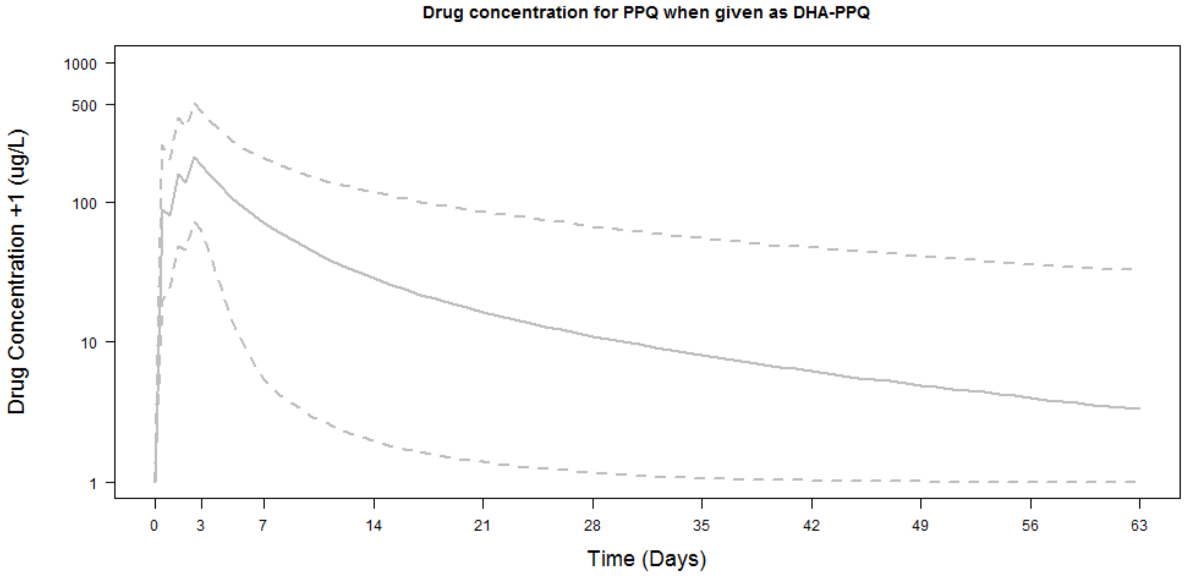


**Figure S1.1** PPQ concentration (in µg/L) over time for a population of 5,000 patients, treated with DHA-PPQ parameterized as in Table S1.1 with drug dosing as in Table S1.2. The solid line is the median population concentration at each day and the dashed lines are the 5% and 95% quantiles. The figure follows patients for 63 days (the maximum length of patient follow-up investigated).

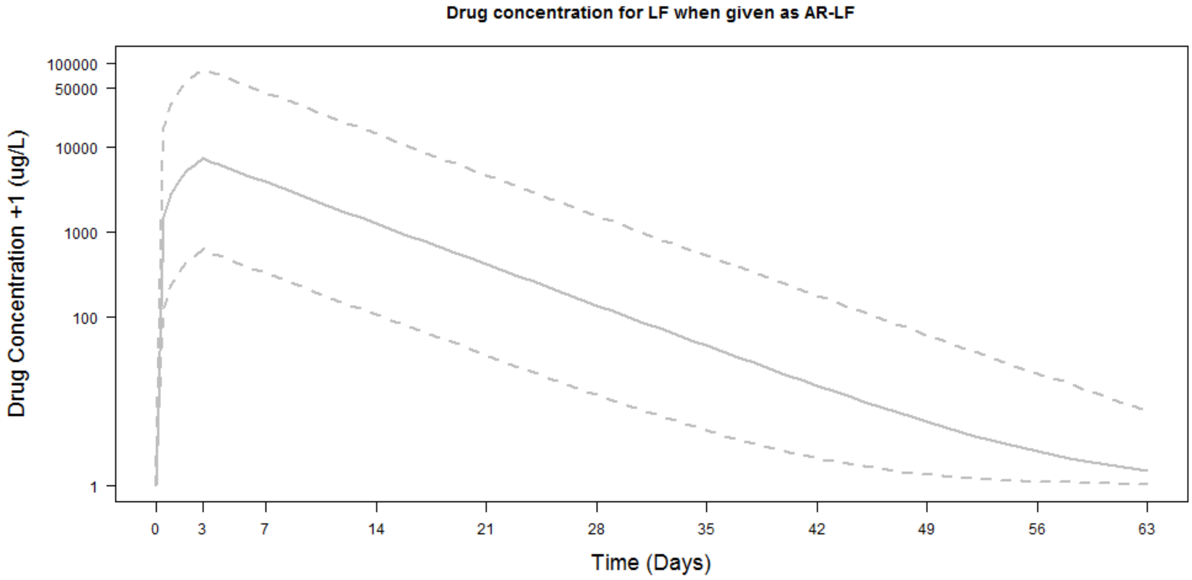


**Figure S1.2** LF concentration (in µg/L) over time for a population of 5,000 patients, treated with AR-LF parameterized as in Table S1.1 with drug dosing as in Table S1.2. The solid line is the median population concentration at each day and the dashed lines are the 5% and 95% quantiles. The figure follows patients for 63 days (the maximum length of patient follow-up investigated).

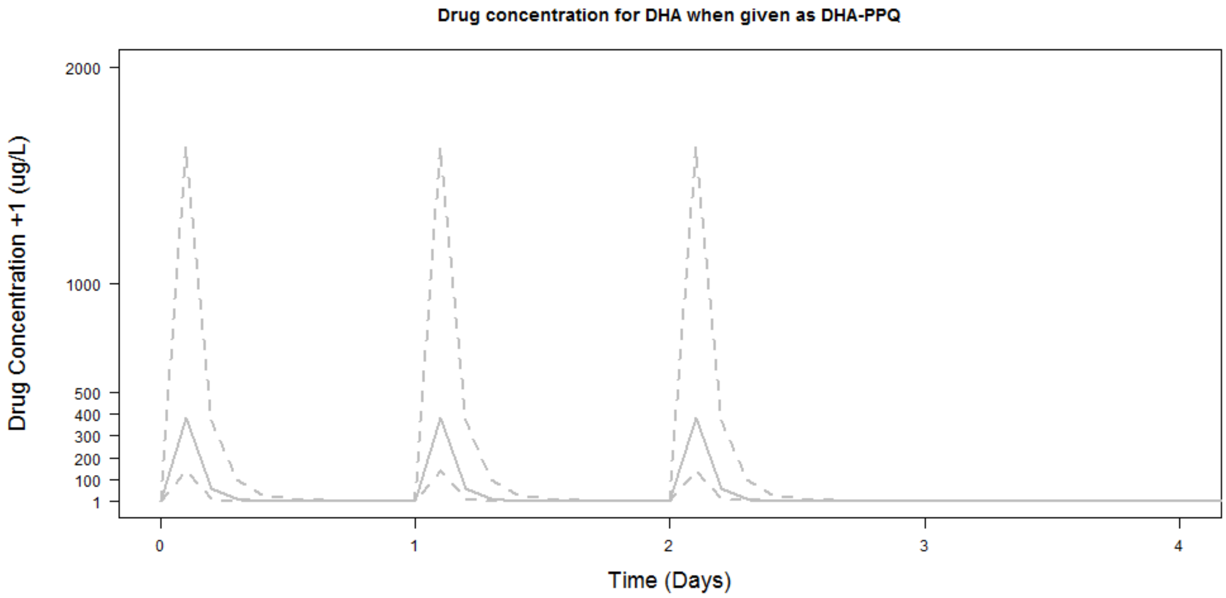


**Figure S1.3** DHA concentration (in µg/L) over time for a population of 5,000 patients, treated with DHA-PPQ parameterized as in Table S1.1 with drug dosing as in Table S1.2. The solid line is the median population concentration at each day and the dashed lines are the 5% and 95% quantiles. The figure follows patients for 4 days (after which all artemisinins have decayed to non-effective and/or zero concentrations).

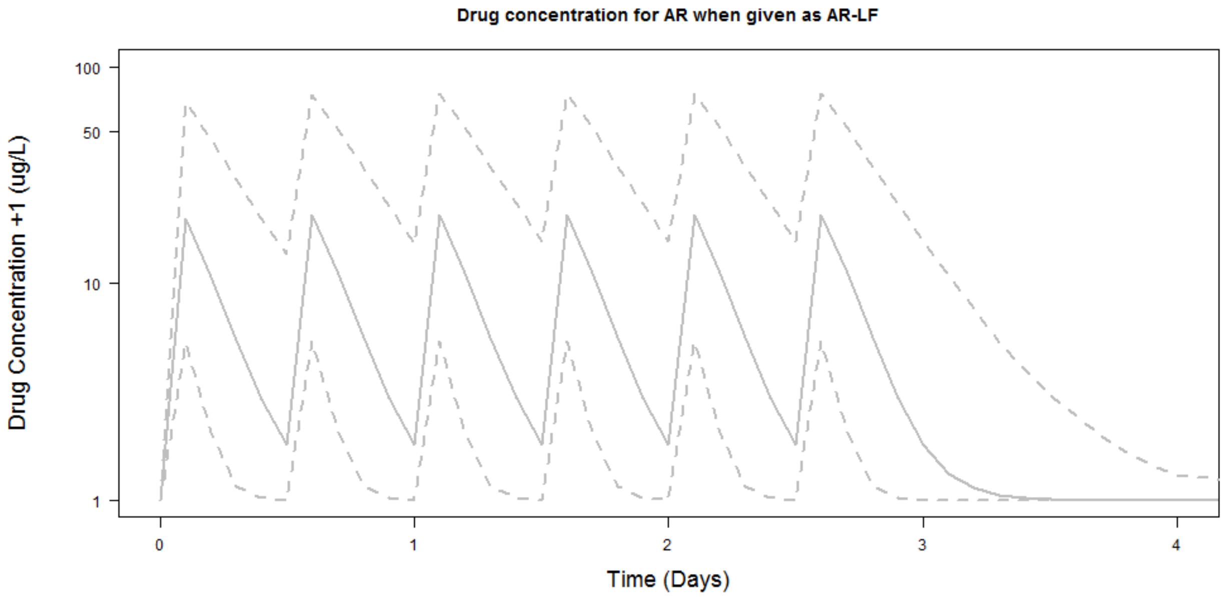


**Figure S1.4** AR concentration (in μg/L) over time for a population of 5,000 patients, treated with AR-LF parameterized as in Table S1.1 with drug dosing as in Table S1.2. The solid line is the median population concentration at each day and the dashed lines are the 5% and 95% quantiles. The figure follows patients for 4 days (after which all artemisinins have decayed to non-effective and/or zero concentrations).

#### Supplementary Material Part 2: Additional Results and discussion

##### 2.1 Additional results obtained when changing model parameters

Four important model parameters were varied to assess their impact on results: The BIC value, the blood sampling limit, the lower limit of the parasite number distribution in the patient sample taken at treatment, the number of *Ampseq* loci genotyped.

###### 2.1.1 Changing the bioinformatics cut-off (BIC) value.

(see Figures S2.1 and S2.2)

The value BIC=1% was used in the baseline calculations (i.e., minority alleles were detected if they exceeded 1% of total reads). This threshold is user-defined by the genotyping software when AmpSeq markers are genotyped (1) so assessing the impact of this parameter on the accuracy of molecular correction is important. As noted by Gruenberg et al., "a stringent cut-off is required for excluding sequencing errors; on the other hand, a less stringent cut-off would be desirable for maximized detection of minority clones". Here, we increased this threshold to 2% to assess the impact of a higher cut-off point. We also reduced it to 0% to assess the difference between 1% and 2% cut-offs from the hypothetical perfect detection of minority clones that would occur at BIC->0%, noting that ->0% is unfeasible in practice because such a cut-off permits the inclusion of sequencing errors and contaminations.

Failure rate estimates for DHA-PPQ and AR-LF with low and high MOI and a range of FOI values are shown in Figure S2.1 for BIC_.0% and Figure S2.2 for BIC=2%. When compared to BIC=1% (Figure 1 of main text), a BIC-> 0% resulted in slightly higher failure rate estimates and BIC=2% resulted in slightly lower failure rate estimates. In both cases, the difference was negligible and failure rate estimates obtained using BICs of ->0%, 1% and 2% were very close. In other words, the currently proposed threshold of BIC=1% has negligible difference to a hypothetical perfect detection scenario. If genotyping errors were to necessitate a higher BIC, our results indicate that BIC=2% could be used with minimal loss of accuracy compared to BIC=1%.

###### 2.1.2 Changing the blood sampling limit

(see Figures S2.3)

The blood sampling limit (the parasitaemia of a clone required for it to be physically included in a finger-prick blood sample used for genotyping) was $10^8$ total parasites in the baseline model. This limit was calculated based on realistic blood sampling processes (see Supplementary Material, Part 1 i.e. section 1.7) but giventhe ability of AmpSeq to detect low frequency alleles, it was necessary to check this assumption was not biasing results. The blood sampling limit was therefore reducing to $10^7$ total parasites (i.e. a lower density clones would be included in the blood sample).

Failure rate estimates obtained with the lower blood sampling limit are shown for DHA-PPQ and AR-LF in Figure S2.3. Results were qualitatively extremely similar to the baseline model (Figure 1 of main text). There was an extremely small increase in failure rate estimates at higher FOI (8 and 16) of ~0.02% when using lower number of matches (≥1 or ≥2) to classify a recrudescence. In short, results were functionally identical to the baseline assumption of a blood sampling limit of $10^8$, so the assumed value of the blood sampling limit did not appear to affect the failure rate estimates obtained using AmpSeq markers.

##### 2.1.3 Changing the lower limit of the initial parasite number distribution

(see Figures S2.4)

The lower limit of the log-uniform distribution was changed from $10^{10}$ parasites (the baseline lower limit) to $10^{8}$ parasites (the upper limit remains as $10^{11}$ parasites). This allows us to investigate whether the accuracy of failure rate estimates generated using AmpSeq markers was affected by assuming a wider range of initial parasitaemia across clones (which will lead to increase proportions of low density clones). The true failure rate changed as the range of this distribution changed: 8.8% and 4.4% for DHA-PPQ in high and low MOI settings respectively and 10% and 6.6% for AR-LF in high and low MOI settings respectively. These true failure rates are slightly lower than the baseline scenarios, presumably because MOI is held constant so more low-density clones are present in the initial infections, and such low-density clones are less likely to recrudesce. However, the absolute change in true failure rate was negligible, so this effect did not appear to be large. The BIC was 1% and blood sampling limit was $10^{8}$, both as for the baseline model.

Failure rate estimates using a wider initial parasite number distribution (i.e. reducing log distribution of initial parasite number down to $10^{7}$) are shown for DHA-PPQ and AR-LF in Figure S2.4. Failure rate estimates were slightly lower in both cases, though this should be considered relative to the slightly lower true failure rate. The difference between each estimate and the true failure rate, and thus the qualitative conclusions, were identical to the baseline model i.e. classifying a recrudescence at ≥2 or 3 matches accurately recovered the true failure rate for both drugs, both MOI settings and all FOI values.

##### 2.1.4 Increasing the number of Ampseq loci

(see Figures S2.5 and S2.6)

We calculated failure rate estimates using 4 AmpSeq markers (including *csp*) and 5 AmpSeq markers (including *csp* and *msp-7*) – this represents inclusion of the less diverse markers in the marker data set. Failure rate estimates obtained using 4 or 5 AmpSeq markers are shown in Figures S2.5 and S2.6 respectively. In both cases, classifying a recrudescence at ≥2 matches no longer produces accurate failure rate estimates (compared to using 3 AmpSeq markers (*cpp*, *cpmp*, *ama1-D3*)) and leads to over-estimation of failure rate at higher FOI. This effect arises because inclusion of less diverse markers increases the probability that reinfections share, by chance, a given number of alleles with clones from the initial infection. There is consequently a slightly increased misclassification of reinfections as recrudescences. Our results indicated that if genotyping were to be done on 4 AmpSeq markers, a recrudescence should be defined as ≥3 matches, and for 5 AmpSeq markers a recrudescence should be defined as ≥4 matches. In summary, genotyping 4 or 5 AmpSeq markers does not significantly increase molecular correction accuracy compared to genotyping using 3 AmpSeq markers. We do note the practical point that genotyping additional markers will be useful in the case that any other markers fail to amplify or are otherwise corrupted in the genotyping process. This suggests that the best threshold is ≥(n-1) where n is the number of markers genotyped.

The AmpSeq markers used in these simulations were selected in highly SNP-polymorphic regions so there is a high number of alleles for each marker. A lower allele diversity has only been observed for *csp* in one geographic area i.e. PNG (with only 3 alleles (1)). Populations with low genetic diversity have not been genotyped using these markers to date. If/when lower diversity data-sets become available, this modelling work can be repeated to quantify the accuracy of failure rate estimates in such areas. The impact of lower genetic diversity would be to increase failure rate estimates due to "identity by chance" i.e. more reinfections will be misclassified as recrudescence due to them sharing alleles purely by chance. In such areas, it may be necessary to consider a) genotyping a larger number of markers and b) to use higher (more stringent) thresholds of matching loci to classify a recurrence as a recrudescence (i.e., the ≥2/3 threshold may over-estimate true failure rate and a =3/3 threshold may be superior). Use of AmpSeq in clinical trials requires genotyping the initial blood samples and quantifying the level of genetic diversity of those samples, such that informed decisions around the total number of markers to analyse and the threshold chosen can be made. Obtaining accurate MOI and allele diversity estimates is possible with AmpSeq due to their high resolution (2). Estimates of FOI (this parameter can be derived from annual entomological inoculation rate (aEIR), see SI of (3)) should also be obtained where possible. This epidemiological information can then be used to optimize AmpSeq analysis for a given clinical trial.

#### 2.2 Additional discussion of results.

##### 2.2.1 Impact of Multiplicity of Infection (MOI) and drug type.

The multiplicity of infection at time of treatment and drug used (DHA-PPQ or AR-LF) appeared to have little impact on the qualitative results (Figure 1 of main text and Figures S2.1 to S2.6 below); this is consistent with results obtained when simulating other types of marker (3, 4).

##### 2.2.2 Impact of Force of Infection (FOI)

The effect of the Force of Infection (FOI) is shown on Figure 1 of main text and Figures S2.1 to S2.6 below.

When FOI is zero, there is no possibility of re-infection and failure rate estimates did not change as the matching threshold was altered (an obvious result because, when FOI=0, all recurrences must be recrudescence). In these circumstances, a slight under-estimate of true failure rate (<1%) occurred in two scenarios:

(iii) If the recrudescence never reaches patency during follow-up i.e. never grows to $>10^8$ total parasites which is the limit of detection by light microscopy, see methodology in (3). This means the patient is erroneously classed as having cleared their initial infection.

(iv) If the recrudescent clone was comparatively low-density clone at treatment so was not detected in the initial sample (thus the recurrence is misclassified as a reinfection).

The under-estimate was larger for DHA-PPQ scenarios than AR-LF – consistent with previous work indicating that some recrudescences occur later than 42 days after treatment with DHA-PPQ, but nearly all have occurred by 28 days following treatment with AR-LF.

As FOI increased, it started to have an impact because introducing reinfections had two consequences:

(i) There was a higher likelihood of a recurrence containing both recrudescent clone(s) and new infections, and thus a higher chance for recrudescent clones to be below the detection threshold in the recurrent sample (i.e., some truly recrudescent alleles may not be observed in the recurrence).

(ii) More clones, and hence alleles, are likely to be present in the recurrent blood sample so the chance of finding a match with the initial infection purely by chance is increased. This would result in a reinfection being misclassified as a recrudescence (noting this chance will be close to 0 when a match is required at all markers). This occurred at all matching thresholds, but misclassification of reinfection with lower thresholds resulted in higher failure rate estimates and is why failure rate estimates are highest with FOI 16 using a matching threshold of ≥1 but lowest with a threshold of =3.

This affect the relative performance of different matching thresholds as described in the next section.

**2.2.3 Choice of matching threshold to define a recrudescence**

Figure 1 of main text shows that the impact of FOI is heterogeneous at the different matching thresholds used to classify a recrudescence, so the key operational question is thus: What matching threshold produced failure rate estimate closest to the true failure rate, and is this threshold robust for multiple drugs and in multiple MOI and FOI settings? The results can be summarised as follows:

- A matching threshold of ≥1 appeared unsuitable because failure rates were severely overestimated at moderate or high FOI values (because there is high chance of a reinfection sharing, by chance, an allele present in the original infection, leading to reinfections being misclassified as recrudescences).
- A matching threshold of ≥2 greatly reduced the probability of matching-by-chance and returned accurate failure rate estimates.
- A matching threshold of =3 generally slightly under-estimated failure rates.

In summary, aside from the high MOI, FOI=16, AR-LF scenario (where a threshold of ≥2 caused a slight over-estimate of true failure rate), using a threshold of ≥2 appeared to be highly robust and returned the most accurate failure rate estimates in all scenarios. This is supported by results obtained when genotyping a larger number of Ampseq (see 2.1.4 above) where results indicated the most robust matching threshold was ≥(n-1) where n is the number of markers genotyped.

**References.**

1. Lerch A, Koepfli C, Hofmann NE, Messerli C, Wilcox S, Kattenberg JH, Betuela I, O'Connor L, Mueller I, Felger I. 2017. Development of amplicon deep sequencing markers and data analysis pipeline for genotyping multi-clonal malaria infections. BMC Genom 18:864.
2. Lerch A, Koepfli C, Hofmann NE, Kattenberg JH, Rosanas-Urgell A, Betuela I, Mueller I, Felger I. 2019. Longitudinal tracking and quantification of individual *Plasmodium falciparum* clones in complex infections. Sci Rep 9:3333.
3. Jones S, Kay K, Hodel EM, Chy S, Mbituyumuremyi A, Uwimana A, Menard D, Felger I, Hastings IM. 2019. Improving methods for analysing anti-malarial drug efficacy trials: molecular correction based on length-polymorphic markers *msp-1*, *msp-2* and *glurp*. Antimicrob Agents Chemother 63:e00590-19.
4. Jones S, Plucinski M, Kay K, Hodel EM, Hastings IM. 2020. A Computer Modelling Approach To Evaluate the Accuracy of Microsatellite Markers for Classification of Recurrent Infections during Routine Monitoring of Antimalarial Drug Efficacy. Antimicrob Agents Chemother 64:e01517-19.

**Figure S2.1.** As for Figure 1 of the main text, but with BIC changed from 1% to ->0%.

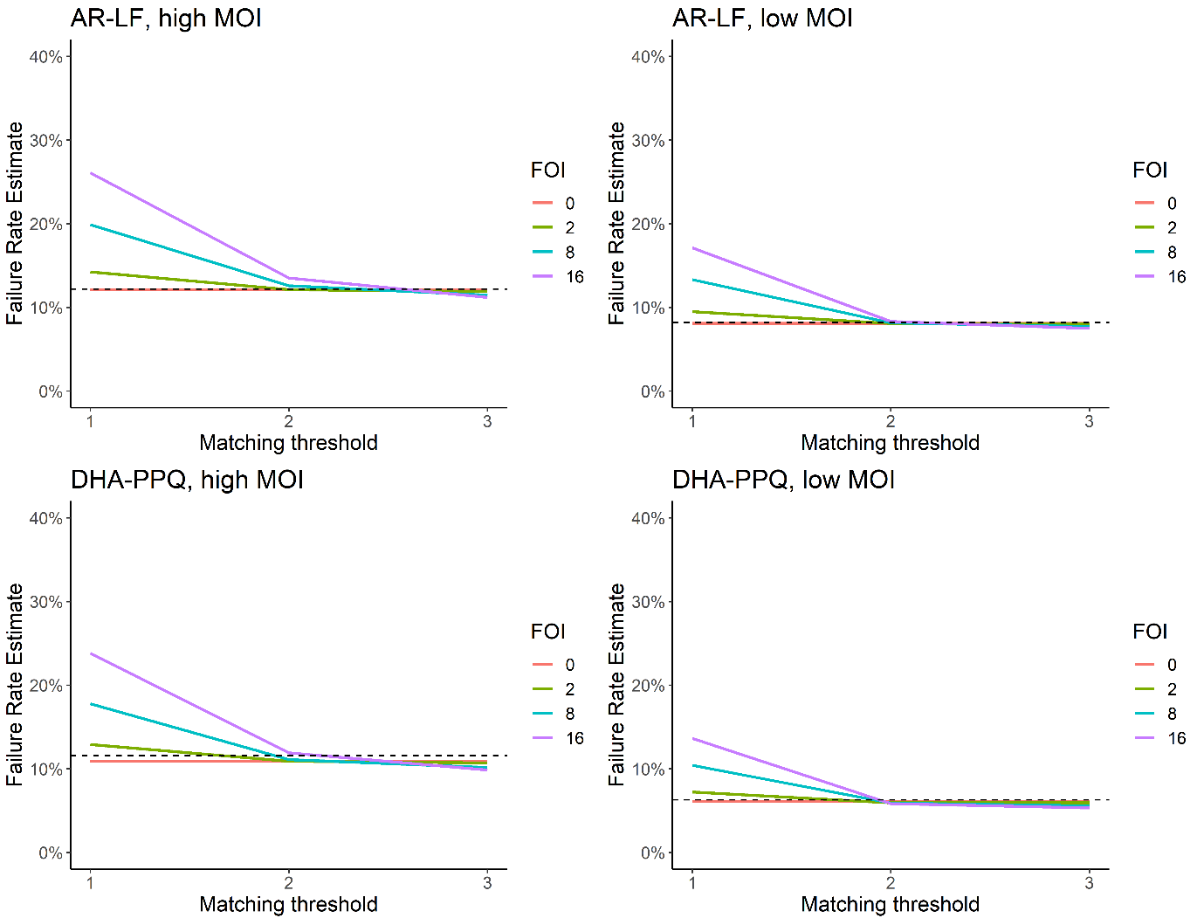

**Figure S2.2.** As for Figure 1 of the main text, but with BIC changed from 1% to 2%.

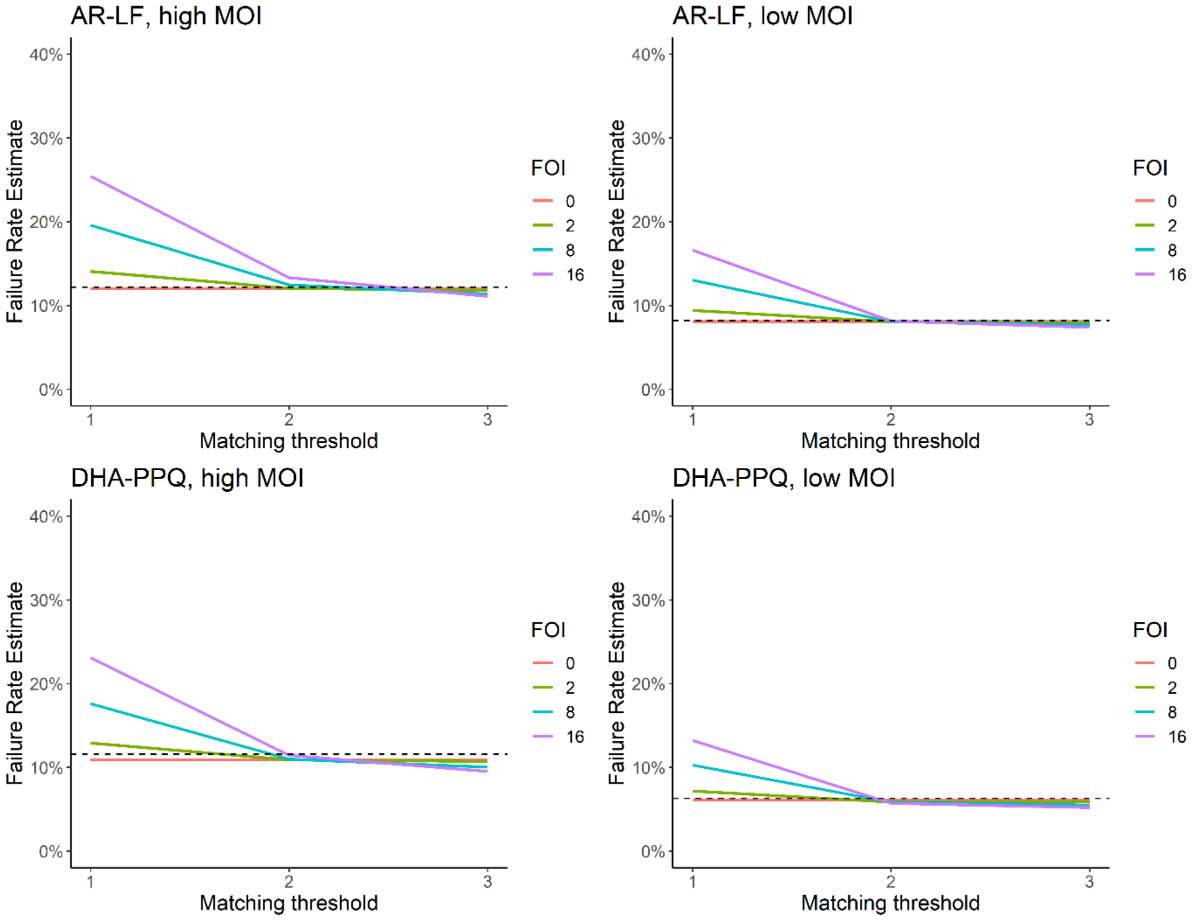

**Figure S2.3.** As for Figure 1 of the main text, but with blood sampling limit changed from $10^8$ to $10^7$.

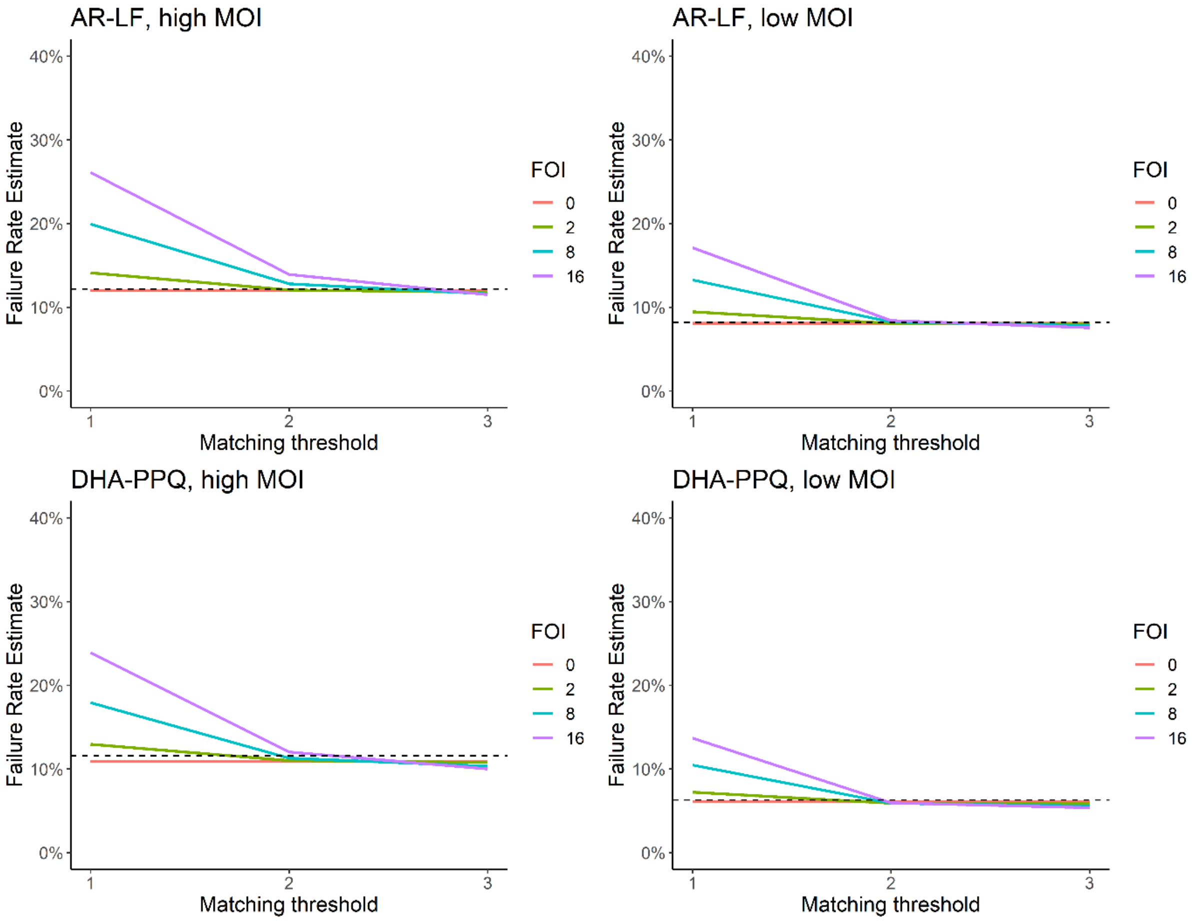

**Figure S2.4.** As for Figure 1 of the main text, but with initial parasite number range changed from $10^{10}$-$10^{11}$ to $10^{8}$ -$10^{11}$.

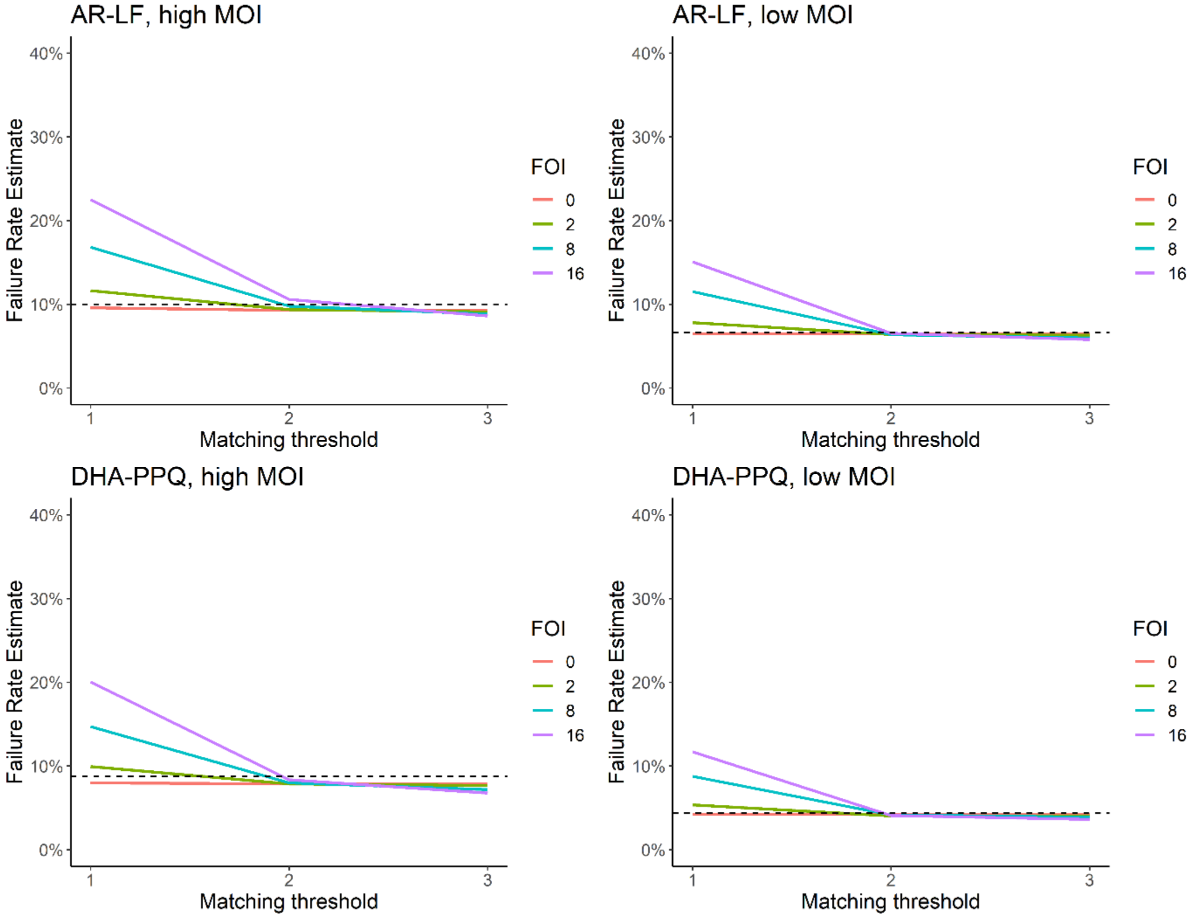

**Figure S2.5.** As for Figure 1 of the main text, but with failure rate estimates calculated using 4 markers (including *csp*)

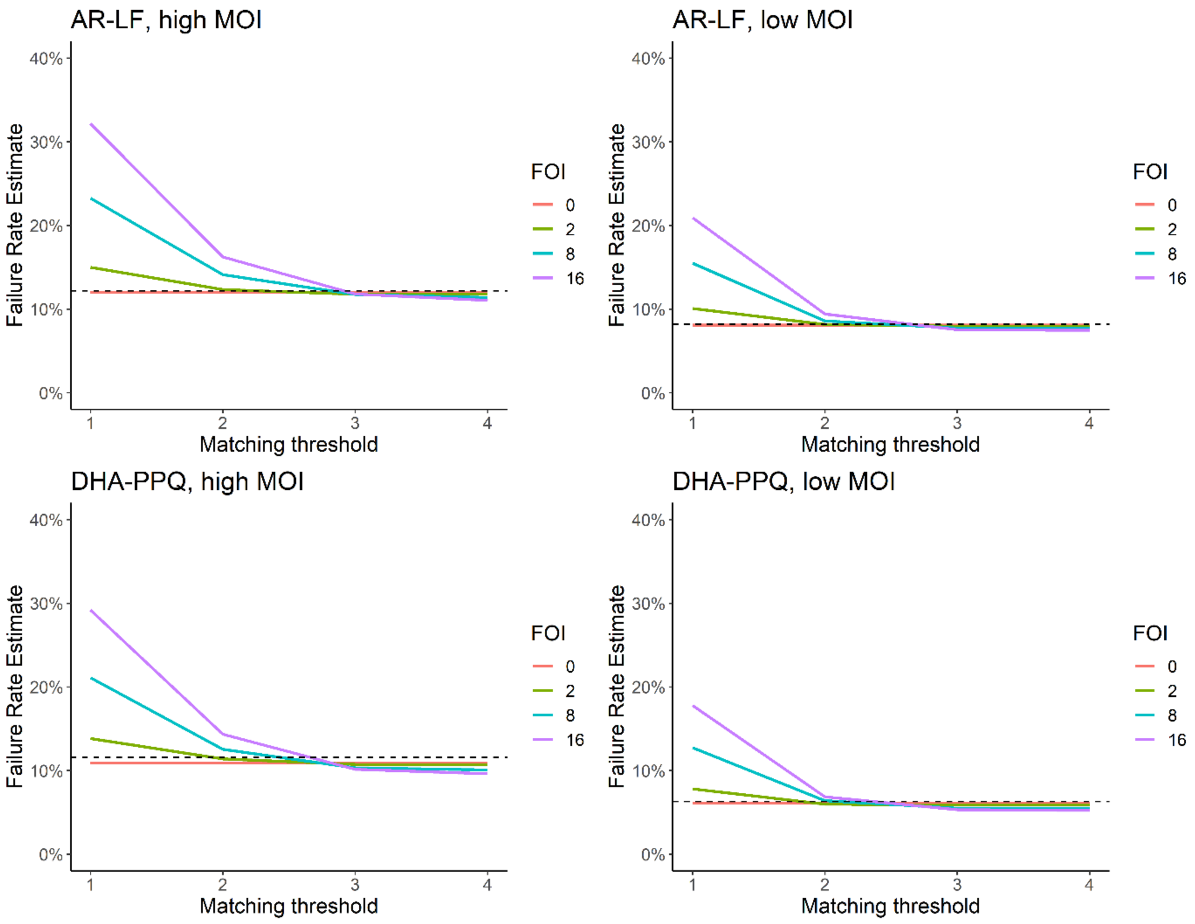

**Figure S2.6.** As for Figure 1 of the main text, but with failure rate estimates calculated using 5 markers (including *csp* and *msp-7*)

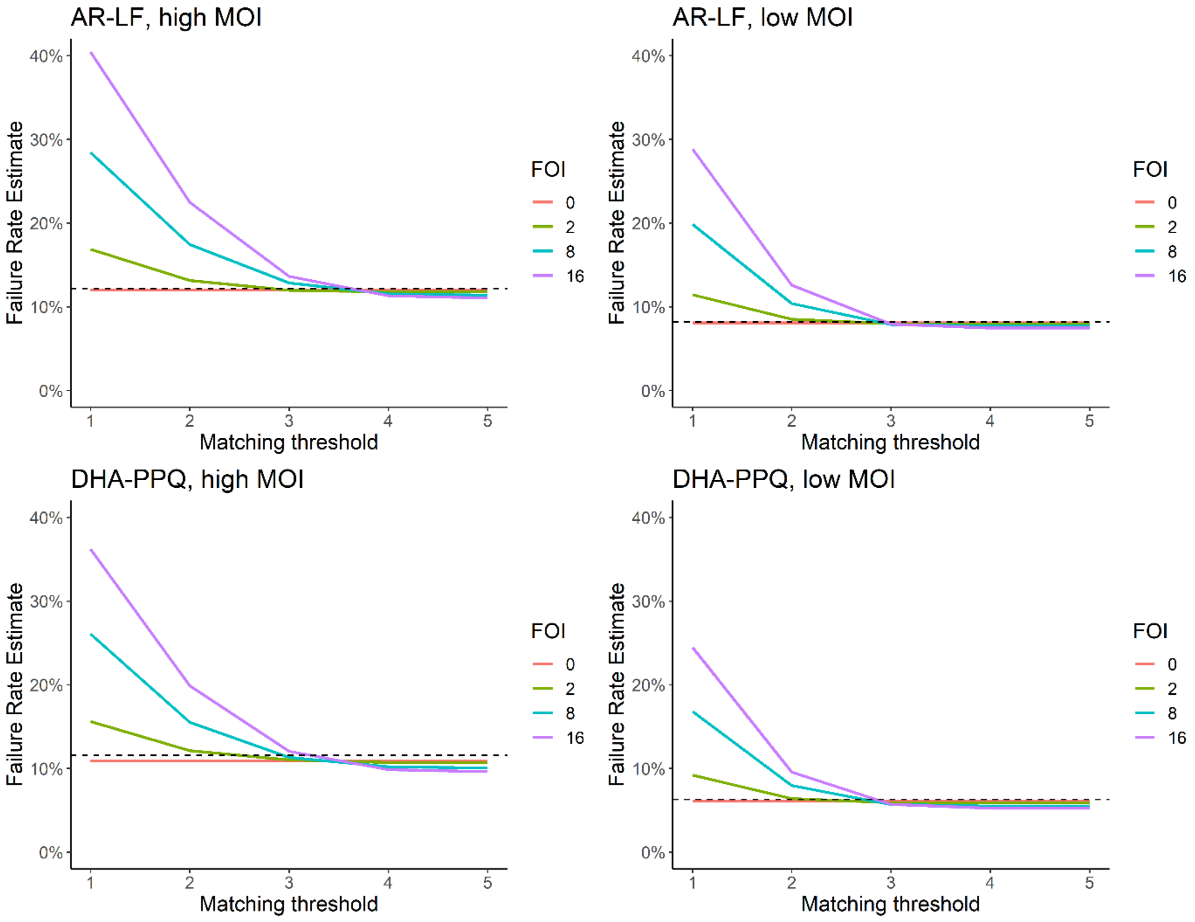

**Supplementary Material Part 3: Simulating the potential impact of gametocytes on the accuracy of molecular correction.**

#### 3.1 Methods: Inclusion of gametocytes in the mechanistic model

The main manuscript focuses on simulation and analysis of antimalarial drug clinical trials. A potential threat to molecular correction in such trials comes from Ampseq genotyping detecting genetic signals arising from gametocytes that are present at treatment but decline relatively slowly post-treatment (see (1) for further discussion in this context). Space limitations in the main text dictate that we provide details of how gametocyte signals were calculated and interpreted asa self-contained section here in the Supplementary Material.

Both asexual and sexual parasite stages of *P. falciparum* may circulate in the blood of an infected host. Asexual merozoites can produce gametocytes that are the transmission stages which subsequently infect the mosquito vector to allow onward transmission of the infection. These gametocytes contain the full complement of genetic material including *msp-1*, *msp-2* and *glurp*, microsatellites, and AmpSeq loci. AmpSeq, in common with other DNA genotyping, cannot distinguish genetic signals from asexual forms and gametocytes, so the presence and detection of gametocyte signals in blood samples may affect molecular correction. The chief concern is that gametocyte alleles persisting from initial infections can be detected and cause later reinfections to be misclassified as recrudescence (i.e. the gametocytes carry signals that will be shared between original and recurrent infections, meaning the latter will be classified as recrudescent). Empirical data show gametocytes can remain detectable in a patient's blood for an average of 55 days (2) after ACT drug treatment. It is therefore important to consider how detection of these gametocyte genetic signals may affect molecular correction. We therefore simulate the dynamics of gametocytes post-treatment to quantifier the likely implications for molecular correction.

None of the current front-line drugs (apart from primaquine) kill mature *P. falciparum* gametocytes. Drugs will, however, determine gametocyte densities post-treatment, both by killing their merozoite 'parents' and by killing some immature stages of gametocyte development. This cuts off the supply of mature gametocytes. Once the supply of mature gametocytes has been cut-off, the gametocytes decline at a rate described by their half-life in the circulation. Changes in gametocyte number over time post-treatment can therefore be tracked given three pieces of information:

(i)The starting number of mature gametocytes at time of treatment:

Each clone of malaria parasites in each infected host is assigned an initial asexual parasitaemia in our calculations. Clones may also contain gametocytes which are generally quantified as a percentage of the total parasitaemia of that clone. Estimates of the starting gametocytaemia vary considerably in the literature; data from Kenya indicate that gametocyte density may be between 1% and 10% of asexual parasitaemia with patient age and symptoms (the presence of fever) affecting this percentage (3). Given the initial asexual parasitaemia and percentage gametocytes allows us to calculate the starting number of mature gametocytes.

(ii) The lag time until gametocyte density starts to decline post-treatment.

Gametocytes take approximately 10 days to develop into mature sexual parasite stages. Antimalarial drugs kill the asexual parasites and hence cut off the supply of gametocytes, so the simple expectation is that the lag time before gametocytaemia starts to decline after treatment would be around 10 days. The lag period can be directly observed *in vivo* and is not 10 days. Most drugs have a lag time of around 5 to 7 days (implying the drugs kill gametocytes in their first 3 to 5 days of maturation). However, artemisinins (and hence ACTs) appear to have a lag time of 3 days (implying artemisinins kill gametocytes in their first 7 days of maturation).

(iii) The rate of gametocyte decline after the lag period.

After the lag-period, the number/denisty of gametocytes decline according to their half-life ($g_{1/2}$). Gametocyte half-life ($g_{1/2}$) and/or their elimination rate, k, can be extracted from published *in vivo* data on gametocyte circulation dynamics (e.g. (2)) and inter-converted using the equation:

$g_{1/2} = 0.693/k$

The results in Figures 2 and 3 of (2) were analysed in this manner to give $g_{1/2}$ estimates of 2.78, 3.27 and 2.15 days based on comparing gametocyte densities on day 3 and 28. The authors of (2) also provide "mean circulation times" of 4.6, 5 and 6.5 days. The mean circulation time is 1/k, so these figures can be converted into $g_{1/2}$ estimates of 3.2, 3.5 and 4.5 days. A mean circulation time of 6.4 days is given in (4); this translates to a $g_{1/2}$ of 4.4 days. A mean $g_{1/2}$ of 2.4 days is reported in (5). Finally, SI Table 3 of (6) allows calculations of $g_{1/2}$ as 6.86, 4.72, 6.12 and 5.77 days when comparing gametocyte numbers on day 7 and day 42. In summary, estimates of $g_{1/2}$ appears to vary between 2.15 and 6.86 days, possibly reflecting differences in host immunity.

The number of gametocytes of a clone present in the blood at time *t* following treatment can thus be expressed by the following equations:

For t≤x (i.e. during the lag period)

$G_t = P_0\gamma$ Equation 3.1

For t>x (i.e. after the lag period)

$G_t = P_0\gamma e^{-k(t-x)}$ Equation 3.2

Where $P_0$ is the initial asexual parasitaemia of a clone, $\gamma$ is the percentage gametocytaemia of that clone, *k* is the gametocyte elimination rate, and *x* is the lag period before gametocyte numbers fall.

Note that we assume the gametocyte density does not change during the lag period. In fact, it may increase slightly as older infections tend to have higher gametocytaemia and there is speculation that some drug treatments may stimulate gametocyte production. We could allow gametocytaemia to increase during the lag period but this putative effect is ignored for simplicity.

These equations allow us to calculate the number of gametocytes from clones present at time of treatment and still circulating on the day of recurrence. Their density at the time of recurrence determines whether their genetic signal will be detected and hence potentially affect the decision to classify the recurrence as a recrudesce or new infection.

**3.2 Main results: gametocyte dynamics post-treatment.**

We did not attempt a full exploration of the impact of gametocytes on molecular correction because the results would be rather obvious: assuming high gametocytaemia and long half-lives would result in high gametocytes signals at recurrence with a potentially large impact on molecular correction. Conversely, assuming low parasitaemia and short half-life would greatly reduce the possible impact. We also found that estimates of half-life appear to vary between 2.15 and 6.86 days, possibly reflecting differences in host immunity (see above). Here we simply illustrate how gametocyte signals start to become detectable with BIC=1% and compare it to the 25% sensitivity of current methods based on gel electrophoresis. The calculation requires a three-stage process

(1) We start by recording the parasitaemias of recurrences that occur at various days post treatment in our simulated clinical trial. These distributions are the box plots in Figure S3.1; note they are identical across each of the four panels.

(2) We then calculate the gametocytaemia persisting from four illustrative clones that were present at time of treatment. We allow initial gametocytaemia at treatment to be $10^8$ or $10^9$; the latter value is high but plausible e.g. a clone present at treatment with asexual biomass of $10^{10}$ with 10% gametocytaemia, or a clone of $10^{11}$ parasites with 1% gametocytaemia. Gametocyte half life may take either of two values i.e. 2.15 days or 6.86 days which represent the extremes of the range of values we extracted from the literature (see above). This enables us to track four illustrative clones whose gametocyte numbers are shown as the red lines on their corresponding four panels of Figure S3.1. Note that the same blood sampling limit occurs as for asexual forms i.e. only clones whose gametocytaemia is above $10^8$ (represented by the horizontal dotted lines in the panels of Figure S3.1) would enter the finger-prick sample and be potentially detectable.

(3) Finally we plot lines equal to gametocytaemia multiplied by 4 or by 100; this is represented by the green and blues lines respectively in Figure S3.1.

This enables us to interpret the panels as follows: *providing gametocytaemia is above the sampling limit ($10^8$; shown by the horizontal dotted line) at time of recurrence*, then each of the four exemplar clones will be:

- Detectable by Ampseq in all recurrences in the boxplots whose parasitaemia lies below the green line (because gametocytes in that clone are present at >1% of total parasitaemia)
- Detectable by standard length polymorphism (e.g. the standard WHO-recommended methods based on *msp*1, *msp*2 and *glurp*) in all recurrences in the boxplots whose parasitaemias lies below the blue line (because gametocytes in that clone are present at >25% of total parasitaemia)

Malaria clones with “low” gametocytaemia at treatment (which is actually still rather high), are likely to have fallen below the blood sampling limit by the first day of follow-up (day 7) so are highly unlikely to be detected when genotyping recurrences irrespective of whether the gametocytes have long or short half-lives (Figure S3.1, panel A and C).

Malaria clones with high gametocytaemia at treatment do have the potential to be detected during follow-up. Those with short half-life may still be above the blood sampling limit on day 7 (Figure S3.1, panel C ) and the clone would be detectable on that day in almost all recurrences genotyped by Ampseq (green line) and in most recurrences when genotyped by length polymorphism. Clones with a long-half remain above the blood sampling limit for up to 21 days (Figure S3.1, Panel D) during which time they will be detectable by Ampseq in almost all recurrences (green line) and often detectable by length-polymorphism genotyping (blue line).

Figure S3.1 show the detectability of 4 exemplar clones. The overall impact of persisting gametocytes on molecular correction will depend on how frequently these different types of clones are present in the trial study site. Studies carried out in low-gametocytaemia patients will almost certainly not be affected by detection of gametocyte signals, while those enrolling patients with high gametocytaemias may have subsequent molecular correction compromised through detection of genetic signals from gametocytes during follow up, particularly if gametocytes have moderate to long half-lives.

**3.3 Discussion: main implications for gametocyte persistence post-treatment.**

Interestingly, it is the blood sampling limit that is most likely to mitigate the risk of detecting gametocyte genetic signals during follow-up  as most clones with low gametocytaemia at treatment and/or short gametocyte half-lives, rapidly fall below the blood sampling limit meaning they are unlikely to be present in the fingerprick blood samples taken at recurrence.

We let readers decide on how likely high gametocytaemia infections are likely to occur in their trial but note that high gametocytaemias are usually associated with long-established infections (implying some degree of protective immunity) while long-lives may well reflect low levels of acquired immunity. Interesting, even methods based on electrophoresis such as the WHO method using msp1, msp2 and glurp, and the CDC method using microsatellites may have problems with persisting gametocytes (Figure S3.1D). The practical difference is that the electrophoresis techniques only detect gametocytes if they constitute more than around 25% of the parasite biomass and these should be directly observable by light microscopy unless the recurrence is extremely low density. In contrast detecting gametocytes at levels down to 1% is extremely arduous and may well be overlooked.

A high gametocytaemia clone may still be detectable at 7 days even if gametocytes have a very short half-life. (Figure S3.1C). There is some debate about whether day 7 recurrences should be subjected to molecular correction or be automatically classified as early drug failures; our results suggest the latter is the safer course (in practice both methods of classification should be used and compared  to see whether they produce different results). The potential threat of misclassification comes from clones that are highly gametocytaemia at treatment and whose gametocytes have a long half-life (Figure S3.1D). The magnitude of this threat depends on how likely such clones are to exist in the trial.

These results, although illustrative, suggest that the increased potential (compared to current methods) of AmpSeq to detect genetic signals from  persisting gametocyte means that AmpSeq should be carefully rolled out for molecular correction with accompanying operational research to check whether gametocyte signals are likely to affect the failure rate estimates. A plausible initial investigation might therefore compare molecular correction results excluding patients whose blood has detectable gametocytaemia at treatment or recurrence using either light microscopy or molecular methods (2), and compare the results to those obtained when all patients are included.

**References.**

**Figure S3.1.** The potential impact of gametocyte genetic signals on molecular correction. The boxplots show total asexual parasitaemia at the time of recurrence for 5,000 patients with new infections treated with DHA-PPQ (with early treatment failures on day 3 excluded). The potential impact of gametocyte genetic signals is demonstrated by modelling the gametocytaemias post-treatment of four illustrative, gametocytaemia clones present at treatment. The red lines show gametocyte number. The green lines is 100x gametocyte number: since we are assuming BIC=1%, any new infections in the boxplots whose asexual parasite number lies below these green lines will potentially have alleles from these gametocytes detectable when using AmpSeq. The blue lines are 4x gametocytaemia: standard WHO genotyping based on gel-electrophoresis has a sensitivity to detect “minor” genetic signals down to around 25% of the total parasitaemia so any new infections in the boxplots whose asexual parasite number lies below these blue lines will potentially have alleles from these gametocytes detectable using the standard WHO methodology. The horizontal dotted line at $10^8$ is the blood sampling limit: when gametocytaemia falls below this level, gametocytes are highly unlikely to enter a standard fingerpick blood sample

(A) An illustrative clone with $10^8$ gametocytes at time of treatment and a gametocyte half-life of 2.15 days

(B) An illustrative clone with $10^8$ gametocytes at time of treatment and a gametocyte half-life of 6.86 days

(C) An illustrative clone with $10^9$ gametocytes at time of treatment and a gametocyte half-life of 2.15 days

(D) An illustrative clone with $10^9$ gametocytes at time of treatment and a gametocyte half-life of 6.86 days

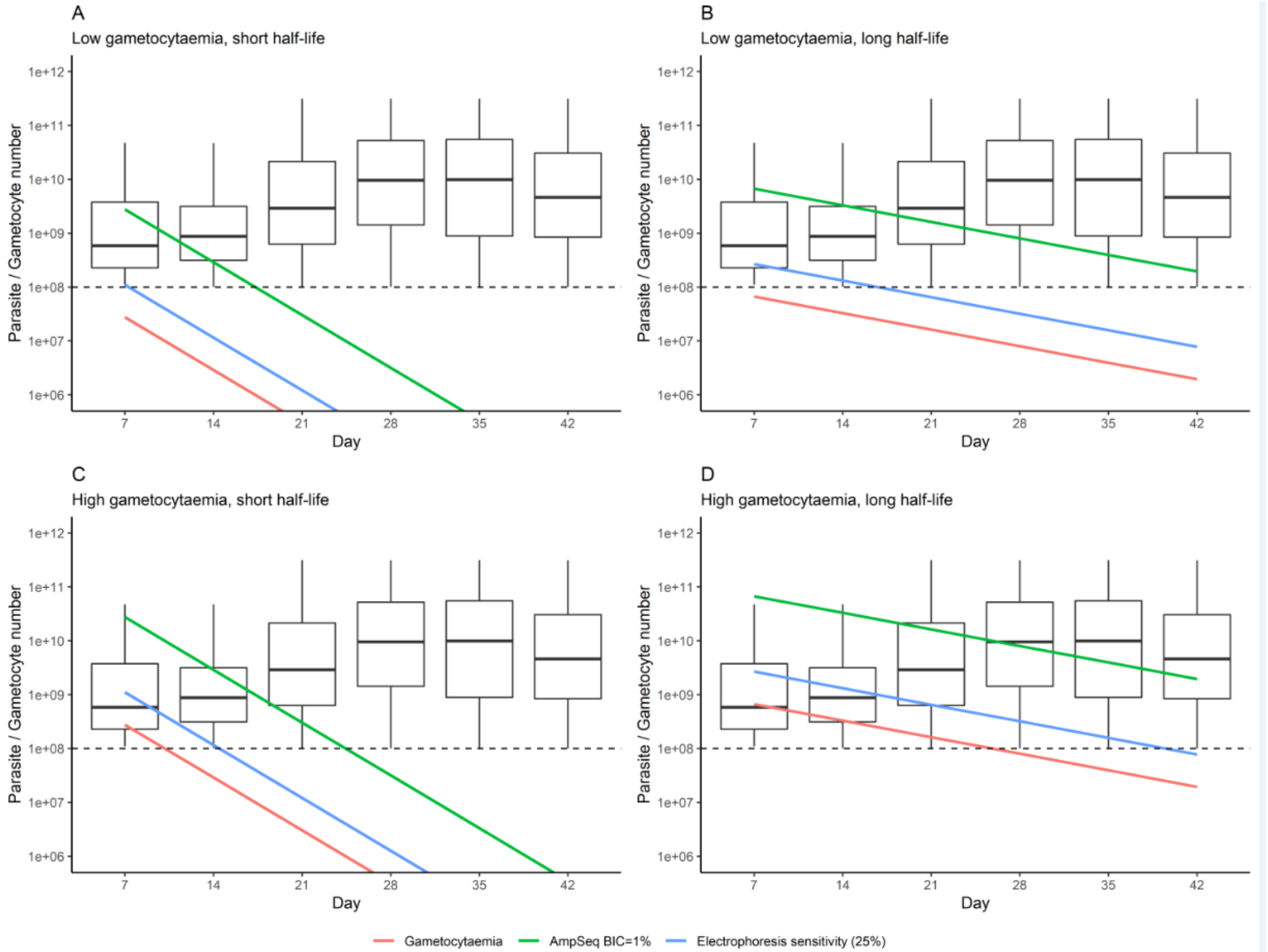
A
Low gametocytaemia, short half-life
B
Low gametocytaemia, long half-life
C
High gametocytaemia, short half-life
D
High gametocytaemia, long half-life
Parasite / Gametocyte number
1e+12
1e+11
1e+10
1e+09
1e+08
1e+07
1e+06
7
14
21
28
35
42
Day
Gametocytaemia
AmpSeq BIC=1%
Electrophoresis sensitivity (25%)
